# Non-invasive brain-machine interface control with artificial intelligence copilots

**DOI:** 10.1101/2024.10.09.615886

**Authors:** Johannes Y. Lee, Sangjoon Lee, Abhishek Mishra, Xu Yan, Brandon McMahan, Brent Gaisford, Charles Kobashigawa, Mike Qu, Chang Xie, Jonathan C. Kao

**Affiliations:** Dept of Electrical and Computer Engineering, University of California, Los Angeles, CA, 90024, United States; Neurosciences Program, University of California, Los Angeles, CA, 90024, United States

## Abstract

Motor brain-machine interfaces (BMIs) decode neural signals to help people with paralysis move and communicate. Even with important advances in the last two decades, BMIs face key obstacles to clinical viability. Invasive BMIs achieve proficient cursor and robotic arm control but require neurosurgery, posing significant risk to patients. Non-invasive BMIs do not have neurosurgical risk, but achieve lower performance, sometimes being prohibitively frustrating to use and preventing widespread adoption. We take a step toward breaking this performance-risk tradeoff by building performant non-invasive BMIs. The critical limitation that bounds decoder performance in non-invasive BMIs is their poor neural signal-to-noise ratio. To overcome this, we contribute (1) a novel EEG decoding approach and (2) artificial intelligence (AI) copilots that infer task goals and aid action completion. We demonstrate that with this “AI-BMI,” in tandem with a new adaptive decoding approach using a convolutional neural network (CNN) and ReFIT-like Kalman filter (KF), healthy users and a paralyzed participant can autonomously and proficiently control computer cursors and robotic arms. Using an AI copilot improves goal acquisition speed by up to 4.3*×* in the standard center-out 8 cursor control task and enables users to control a robotic arm to perform the sequential pick-and-place task, moving 4 randomly placed blocks to 4 randomly chosen locations. As AI copilots improve, this approach may result in clinically viable non-invasive AI-BMIs.

## Main

The only control source in traditional motor BMIs, including computer cursor^1–3^ and robotic arm^4–7^ control, are decoded neural signals (Figure 1a). In intracortical BMIs, recording spikes that have high signal-to-noise ratio (SNR), this traditional approach has produced high-performance cursor control, communication, and robotic arm control^2–7^. However, when neural signals have a low SNR, traditional BMIs also achieve low performance. This particularly impacts non-invasive BMI paradigms, including those decoding from electroencephalography (EEG). Non-invasive BMIs therefore perform significantly worse and have not been widely adopted, even though they present minimal clinical risk.

**Figure 1.**
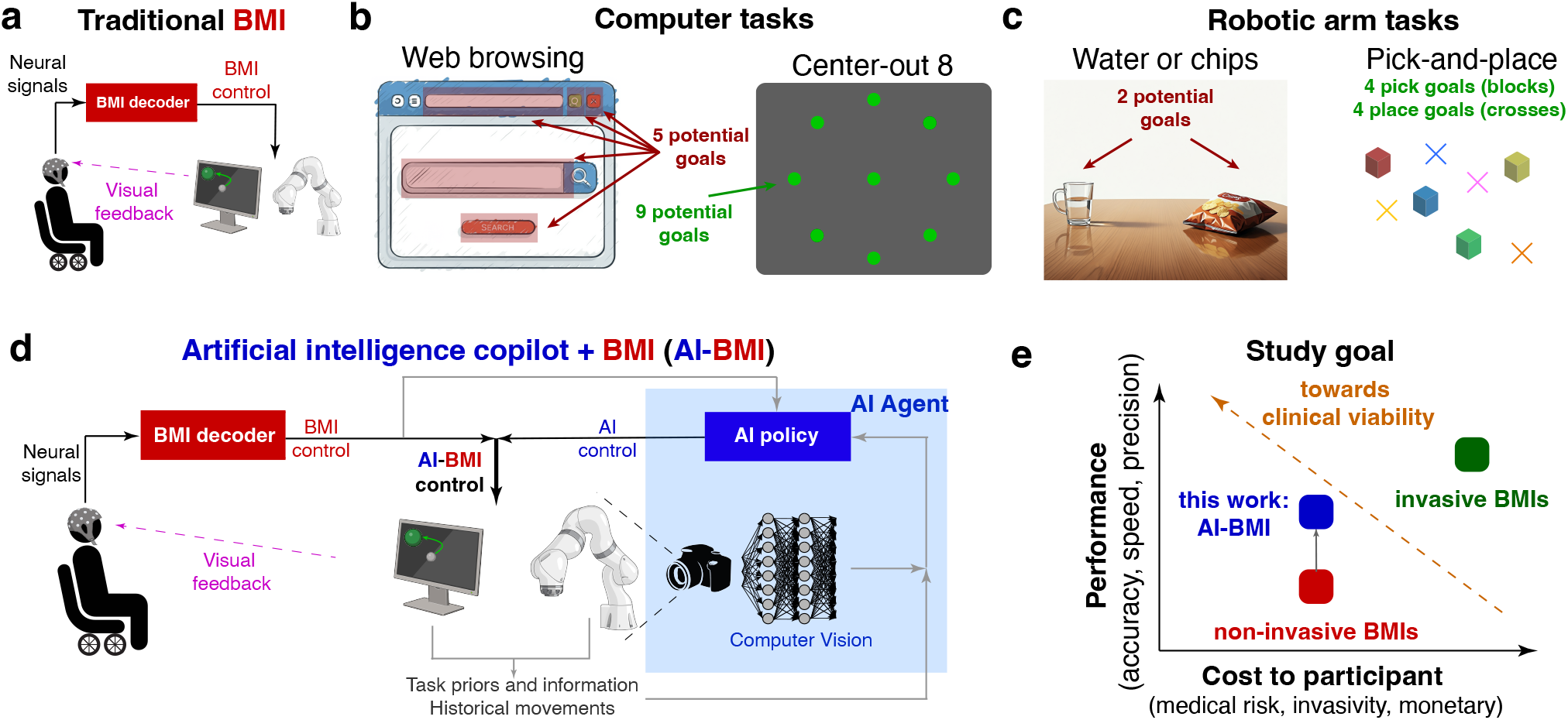
**a**, In traditional brain-machine interfaces (BMIs), neural signals are decoded to control an end effector (e.g., computer cursor or robotic arm). **b**, In computer tasks, there can be a finite number of selectable targets. In web browsing, goals include editing the URL, changing the web page, closing the webpage, entering a search query, or hitting the search button. These goals could be inferred through computer vision (CV). In a BMI benchmark, the center-out 8 task, there are 9 potential targets. **c**, In robotic arm tasks, finite goals often correspond to physical objects. An example may be choosing to drink water or grabbing a bag of chips, both of which could be inferred through CV. In a BMI benchmark, the goals could be pick locations (block) and place locations (crosses). **d**, In an AI-BMI, the BMI and AI agent are combined to produce task-informed actions. The AI uses information, including task priors, past movements, CV, and neurally decoded signals, to infer potential goals and movements. The AI-BMI then helps to complete the task. **e**, The goal of the AI-BMI is to increase task performance without additional cost to participants.

To build performant non-invasive BMIs, we leverage a critical insight: many tasks we perform are *goal-oriented*, where our movements are typically directed toward target locations, such as search bars, buttons, and icons on a computer (Figure 1b), or physical objects such as cups, chips, door handles, keys on a table, or blocks (Figure 1c). In these cases, knowing the user’s goal, a deliberation between a finite number of possible goals, largely determines the movement. Once a goal is known, human actions are often stereotyped and could be aided via an artificial intelligence (AI) copilot. But how do we infer the user’s goal? We reason that, in addition to neural signals, there are other information sources, including task structure, historical movements, and computer vision (CV), that can be used to infer a user’s goal and subsequently aid their movements (Figure 1d).

For example, consider taking a drink from a cup on a table. An intracortical BMI, decoding spikes from populations of neurons, can control the 3D endpoint velocity of a robotic arm to produce a trajectory towards the cup, grasp it, and bring it to the user to drink^4^. Achieving this high-resolution trajectory with only non-invasive EEG signals is very challenging. But an AI copilot, with (1) a camera that sees the cup and (2) task structure knowledge that humans typically grab cups to drink with their mouths, can then process a (3) high-level motor command (such as “move forward”) to infer the user wants to drink from the cup. The copilot would then help the user grab the cup with the appropriate force and bring it to the user to drink. We call this kind of architecture an AI-BMI, illustrated in Figure 1d. Similar kinds of reasoning could be made to infer goals for a computer task. For example, when using a search engine, one’s goal would likely be to select “search” after entering a query into a search bar (Figure 1b), or for typing, the past context gives strong information over the next character selection^8–10^. What if a task doesn’t have a structured goal (such as free drawing with a computer cursor)? In this case, the AI-BMI simplifies to a traditional BMI. But we emphasize that in tasks with goal-directed movements, an AI-BMI can and should use additional contextual and task-specific information to outperform a traditional BMI.

Although an AI copilot could improve both an invasive and non-invasive BMI, we focus on the latter because non-invasive BMIs present less medical risk and cost. Our goal is to build a non-invasive AI-BMI that enables a paralyzed participant to proficiently control computer cursors and robotic arms (Figure 1e). Toward this end, we summarily make two key contributions. First, we develop a novel EEG decoding architecture that uses a CNN’s nonlinear features as the observations of a ReFIT-like Kalman filter (KF)^11^, enabling online closed-loop decoder adaptation (CLDA)^12–14^ (Figure 2a). We characterize the performance of this new decoder architecture and show it can achieve stable performance across days. Second, we demonstrate two copilots, one for cursor control trained with deep reinforcement learning (RL) (Figure 2b) and another for robotic arm control with concurrent computer vision (CV) to automatically detect and grasp or place objects (Figure 2c). We demonstrate AI copilots significantly increase performance in a computer cursor and robotic arm task. We anticipate AI-BMI performance will continue to improve with advancing AI copilots, providing a complementary axis to improve BMIs beyond more traditional approaches of decoder design^11,15–19^ and neural adaptation^12,13,20^.

**Figure 2.**
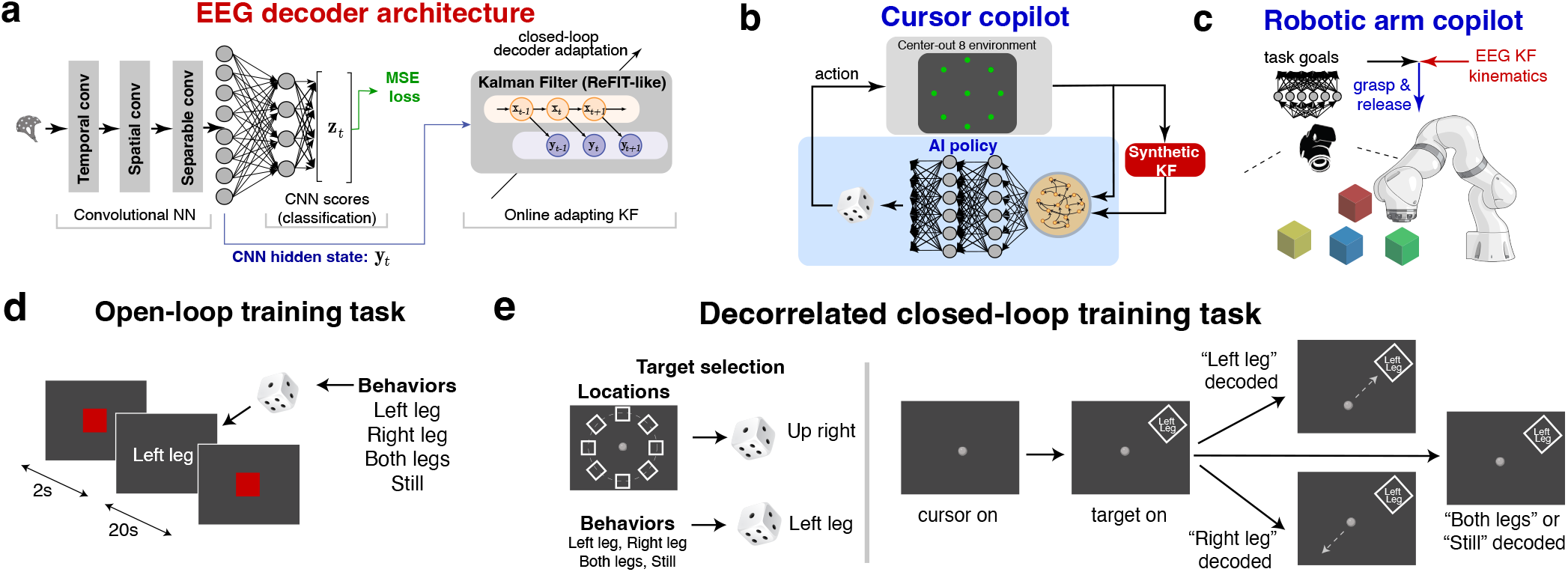
**a**, CNN-KF architecture. The CNN uses the EEGNet, whose final hidden state is the observation of a ReFIT-like Kalman Filter. The Kalman Filter is updated using CLDA. **b**, The cursor copilot is trained with deep reinforcement learning (RL) and synthetic Kalman Filter (KF) output velocities to drive cursor movements towards one of the potential center-out goals. **c**, The robotic arm copilot detects pick and place locations with computer vision (CV). The copilot has a simple policy: when close to a block, it picks it up, and when close to a cross, it places any held object. **d**, Open-loop training task. Text corresponding to one for 4 actions is presented for participants to execute (healthy) or attempt (paralyzed). The order of the 4 actions is randomized. **e**, Decorrelated closed-loop training task. Each trial, one of the 4 actions is placed at any of 8 radial locations. The 2D decoded cursor is projected onto the 1D line connecting the center to the target. The correctly decoded action moves the cursor towards the target, while the opposing decoded action moves the cursor away from the target. The target is presented for 2s and the cursor movement period is 5s.

### A novel non-linear and adaptive EEG decoder

We performed experiments with three healthy (H1, H2, H4) and one spinal cord injury participant (S2) with T5 complete paraplegia, having no movement in his legs. All experiments were approved by the UCLA IRB. Our first goal is to demonstrate stable control of a computer cursor across days. We evaluated performance on an intracortical BMI cursor control benchmark, the center-out task with 8 targets and a 500 ms hold time^2,11,14^. To achieve continuous control, we trained decoders to translate EEG activity corresponding to four discrete behaviors into continuous cursor velocities. Our decoding approach combines deep learning with traditional closed-loop decoder adaptation (CLDA) approaches from invasive BMIs. The decoder was a CNN (the EEGNet^21^) followed by a velocity KF performing ReFIT-like corrections^11,15^, which we call a “CNN-KF” (Figure 2a).

Our design was guided by two principles. First, deep learning architectures empirically achieve higher performance in both offline^21^ and closed-loop experiments^22^. We incorporated an empirically successful CNN adapted from EEGNet^21^. Second, we found that adaptation was important for stable EEG BMI performance. We found EEG occasionally exhibited non-stationarities. For example, it was possible to decode whether EEG activity was recorded in the first or second half a session (Extended Data Figure 2) and as we further characterize below, biases frequently arose in fixed decoders. We therefore incorporated CLDA to stabilize performance. Neural networks are not straightforward to adapt in closed-loop experiments because noisy gradient descent steps are not guaranteed to improve the decoder. We instead froze the CNN, and used its nonlinear features as the observations of a linear KF. The decoded state of the KF was cursor (*x, y*) velocity. We updated the KF parameters in CLDA using the parameter sufficient statistics^12–14^ after ReFIT-like innovations^11^. This enabled us to (1) perform nonlinear decoding while (2) adapting the decoder in real-time to the most recent data. Please see the Methods for additional details on this decoder architecture, training, and CLDA.

To train the CNN-KF, participants first performed an open-loop training task. Participants were prompted with one randomly chosen action out of four total actions (Figure 2d). These four actions corresponded to left, right, up, and down movement classes, where the respective corresponding actions were: left hand, right hand, both hands, feet (H1, H2, H4) and left leg, right leg, both legs, and still (S2). We then trained an initial “seed decoder” from this data (Extended Data Figure 1). This was not the final decoder because we found CNNs may decode features related to eye movement. In closed-loop control, eye movements are highly correlated with cursor and/or target location, meaning that unless the CNN features learn to ignore eye movement activity, the decoder may adapt to decode eye movement features, resulting in spuriously high performance. For example, when the cursor moves from the center to the left-positioned target, the user performs a left motor action while looking to the left of the screen. To ensure we did not decode eye movements, we performed a second training task where presented targets and kinematics were decoupled from motor intent (“decorrelated closed-loop training task”, Figure 2e, see Methods).

This training task was closed-loop^23–26^, where participants controlled a computer cursor using the “seed decoder.” Normally, to move the cursor left, the paralyzed participant would attempt a left leg action. To decorrelate these, we prompted motor intent (“left leg”) to a random target position (“up right” in Figure 2e). When the participant performed the correct motor action (left leg), the cursor moved towards the target (up right). We then trained a CNN on this dataset to decode motor intent. We hypothesized that CNN activity (the hidden state, **y**_*t*_, in Figure 2a, which is the input to the KF) would not reflect any neural features related to eye movements, since eye movements were uninformative of motor intent. To demonstrate this, we performed two analyses. First, we found that the presented target location, reflecting where the eyes looked towards and not motor intent, could not be decoded above chance levels from CNN hidden state (average validation accuracy: 8.0%, chance: 12.5%, Extended Data Figure 3). Second, we used an eye tracker to measure eye position and trained a decoder to predict eye position from the CNN hidden state. This eye position decoder achieved a negative average coefficient of determination on test data across all participants, meaning it could not predict eye position from the CNN hidden state better than the mean of the test data (average test *R*^2^ = *−*0.024, Extended Data Figure 4). Together, these prove that the CNN hidden state, which was the input to the KF, did not represent eye movements. Because the “CNN-KF’ freezes the CNN and only adapts the KF, this architecture and training paradigm yields a nonlinear, adaptable architecture that decodes movement intent and is robust to eye movement artifact signals.

### Stable CNN-KF performance across days

We evaluated the stability and performance of the CNN-KF. Participants performed the center-out task on 5 separate days, distributed over anywhere from 9 to 36 calendar days (see Extended Data Table 1). Similar to Silversmith *et al*.^14^, we initially evaluated center-out 8 performance with large targets that were progressively shrunk after the participant achieved good performance. Circular targets were positioned 8.2 cm from the center target, and initial target diameters were 7 cm, meaning the target could be acquired moving a minimum distance of 4.7 cm. Participants needed to hold the center of the cursor over the target for 500 ms to successfully acquire it. When the participant acquired targets at a rate higher than 10 targets per minute at over 90% success rate, we increased task difficulty by decreasing the target diameter (4.7 cm, 3.5 cm, and 2.9 cm). All participants reached the 3.5 cm target size. Overall performance occasionally decreased when target sizes were decreased. Nevertheless, our goal with these experiments was to demonstrate stable performance over weeks rather than maximal performance.

All participants achieved stable performance across 5 separate sessions (Figure 3a, b). Two key metrics of performance in the center-out 8 task are the percentage of successfully acquired radial targets and the average time to acquire each radial target. We found that, even as the target size decreased, these metrics were stable across all 5 experimental sessions. Participants were always able to complete the center-out 8 task at a high success rate (median: 88% for S2, 100% for all healthy participants, Figure 3d). Trial time expectedly varied based on target size, since smaller targets are more challenging to successfully hold over. Across all sessions, the median trial time was 10.1 s for S2 and between 7.0 s to 9.6 s for healthy participants (Figure 3d). Finally, when summarizing a metric called Fitts’ ITR that factors in target size, success rate, and acquisition time, we did not observe that performance significantly increased from the first day to the last day, but did remain stable. It is possible performance could increase with longer-term CLDA^14^.

**Figure 3.**
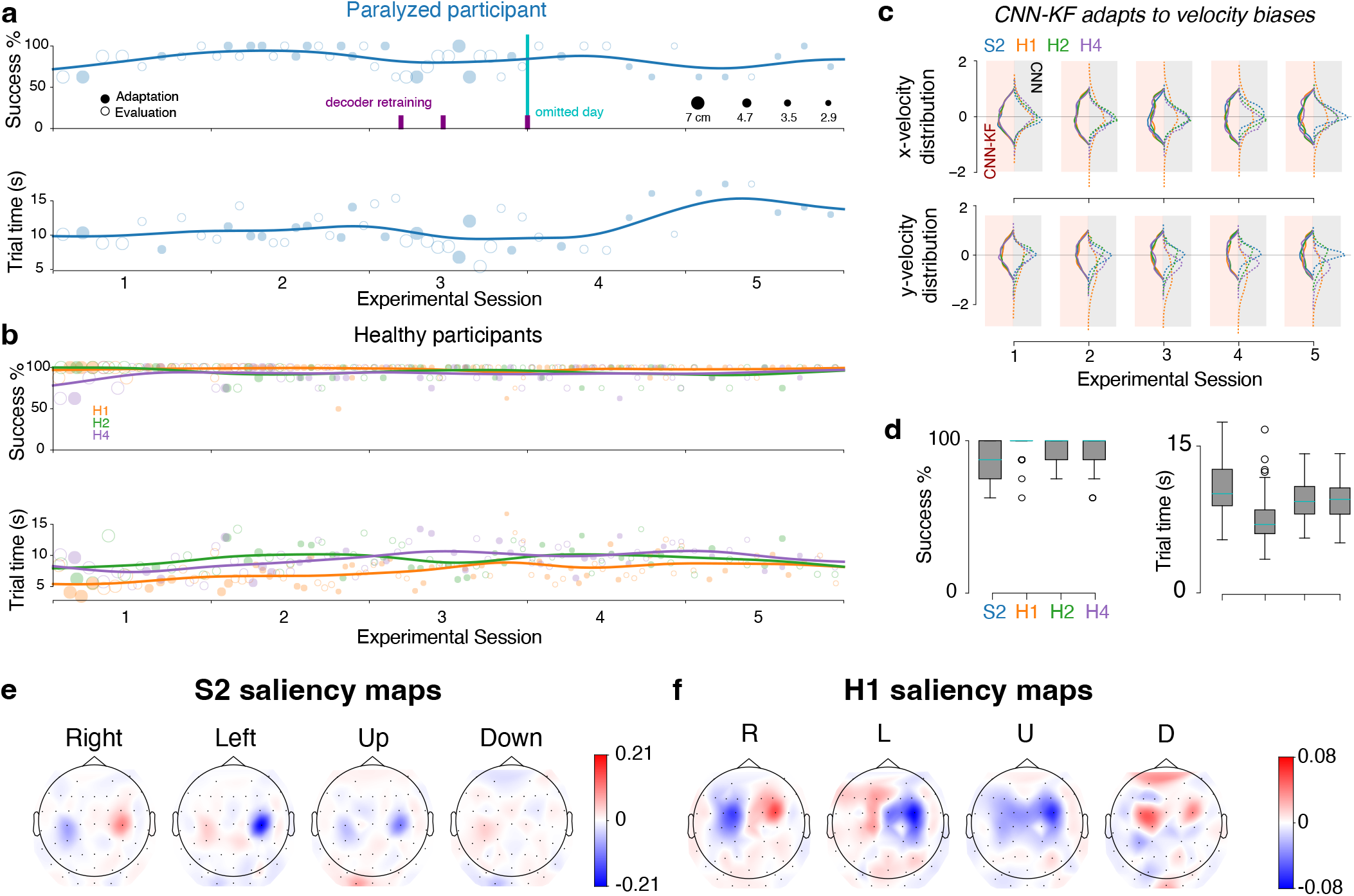
Performance of CNN-KF for center-out 8 task. **a**, Success percentage and trial time for participant S2 on 5 separate sessions while performing the center-out 8 task. The Kalman Filter (KF) statistics of the CNN-KF were incremented during all trials, but were only used to update decoding parameters before and after each adaptation trial. Only trials from center to radial targets are included, and each set of 8 trials is compiled to a single data point. Lines show Gaussian-smoothed averages. Decoder retraining and 1 omitted day due to lower performance (when participant felt paresthesia, see Methods) are indicated in magenta and cyan, respectively. **b**, Same as a, but for three healthy participants. **c**, Half-sided violin plots showing the distribution of *x* and *y* decoded velocity on each day of center-out 8 task for CNN-KF (solid) and CNN only (dashed line). The CNN-KF adapts to velocity biases that emerge in the CNN over days. **d**, Success percentage and trial time across all days. The median of each box plot is shown in cyan. **e**, Saliency maps for paralyzed participant S2. Different hemispheric patterns are activated for each of 4 classes. **f**, Saliency maps for healthy participant H1.

All healthy participants achieved plug-and-play performance, where the same decoder could be adapted day-after-day without recalibration on the decorrelated closed-loop training task (Figure 3b). For S2 (Figure 3a), we decided to retrain the decoder on experimental session 3 (which followed a gap of 8 calendar days) and session 4 (following a gap of 6 calendar days), although we also observed that the decoder did not need to be retrained for session 5 (gap of 19 calendar days). Between experimental sessions 3 and 4, we omitted one session where participant S2 experienced very strong phantom pain. On this day, we were unable to train a high-performance decoder and S2 was unable to perform a full experimental session. One contribution to stable decoding across days was that CLDA on the KF helped to remove biases in decoded velocity due to drifting EEG activity. We show the distribution of decoded velocities when using CNN vs CNN-KF across all participants and experimental sessions (Figure 3c). We observed significant drift in these distributions for some participants with the CNN (dotted lines) but the CNN-KF velocity distribution was relatively unbiased across sessions. Together, these results demonstrate that our decoder approach can yield stable performance across days, and that performance can even be sustained in the same decoder through adaptation, although CNN recalibration may sometimes be required.

What neural features guided proficient decoding? To answer this question, we computed spatial saliency maps that exhibit which EEG electrodes contributed most for each motor class. These saliency maps are based on computing the gradient of the cursor’s velocity update with respect to the magnitude of each electrode’s activity (see Methods). In EEG activity, movement leads to event-related desynchronization, resulting in lower alpha and beta power in contralateral motor areas during movement^27^. Active regions therefore tend to demonstrate a decrease in EEG activity. We found that, as a result of the chosen action sets, hemispheric activity near the motor cortex largely drove cursor movements. The left hemisphere was more active during right movements, the right hemisphere was more active during left movements, both hemispheres were active during up movements, and both hemispheres were less active during down movements (Figure 3e, f, showing saliency maps for S2 and H1, respectively). Together, these results demonstrate that the modulation of hemispheric activity can support proficient and stable 2D EEG cursor control.

### A cursor control copilot more than doubles target hit rates in center-out 8

We hypothesized that an AI copilot that helps infer user goals could increase task performance. In the center-out 8 task, there were only 9 possible goals, corresponding to 9 potential target locations. This imitates, in spirit, how in many applications, there are often only a handful of selectable goals (such as buttons on a computer screen, Figure 1b or likely keys on a keyboard^28^). Our goal was to build a copilot that can infer the user’s goal and help acquire it. Although the possible goals were known a priori in the center-out 8 task due to the task structure, for tasks such as using a computer browser, the possible goal locations can be inferred via computer vision (CV).

We trained a deep RL copilot with proximal policy optimization^29^ (PPO) to assist the user in performing the task. At the highest-level, this copilot observed the BMI decoder trajectories and velocities, and used these inputs to infer what goal the user was trying to acquire, aiding movement towards that goal. The copilot input was the CNN-KF decoded velocities and cursor positions throughout the trial (see Methods). A key advantage of using CNN-KF decoded velocities, and not the high-dimensional EEG, was that this enabled us to train the copilot in a simulated environment where we generated synthetic CNN-KF velocities, removing the need for large experimental datasets. The copilot outputted a probability over potential goals, and affected the cursor velocity through a charge field proportional to the probability^28^. The copilot therefore influenced the velocity of the cursor at every timestep through charge allocation. We trained this copilot in a custom environment where the copilot received positive rewards for movement toward the correct goal and negative rewards for movement away from the correct goal (Methods, Extended Data Figure 5).

We performed our most challenging task condition, where targets were 2.3 cm in diameter with a 500 ms hold time, a more challenging task condition than some intracortical studies^11^. Although the copilot was trained in a simulated environment, we found it generalized to increase closed-loop BMI center-out 8 task performance. Distance-to-target plots^11^ averaged across all single trials summarize the performance improvement (Figure 4a). The cursor copilot reduced the average distance to target significantly faster than without the copilot. For example, in S2, the AI-BMI achieved the same average distance-to-target as the BMI at end of trial (20s) within only 5s. Ultimately, this had the effect of significantly reducing the time it took to complete a trial (Figure 4b). The copilot additionally increased the success percentage of correct target acquisition. S2 and all healthy participants achieved a median accuracy of 100% target success rate (Figure 4c). Together, the increased success rate and decreased trial time means the cursor copilot increased the goal acquisition rate of the task by a factor of 2.26*×* in healthy participants and 4.33*×* in S2 (Figure 4d).

**Figure 4.**
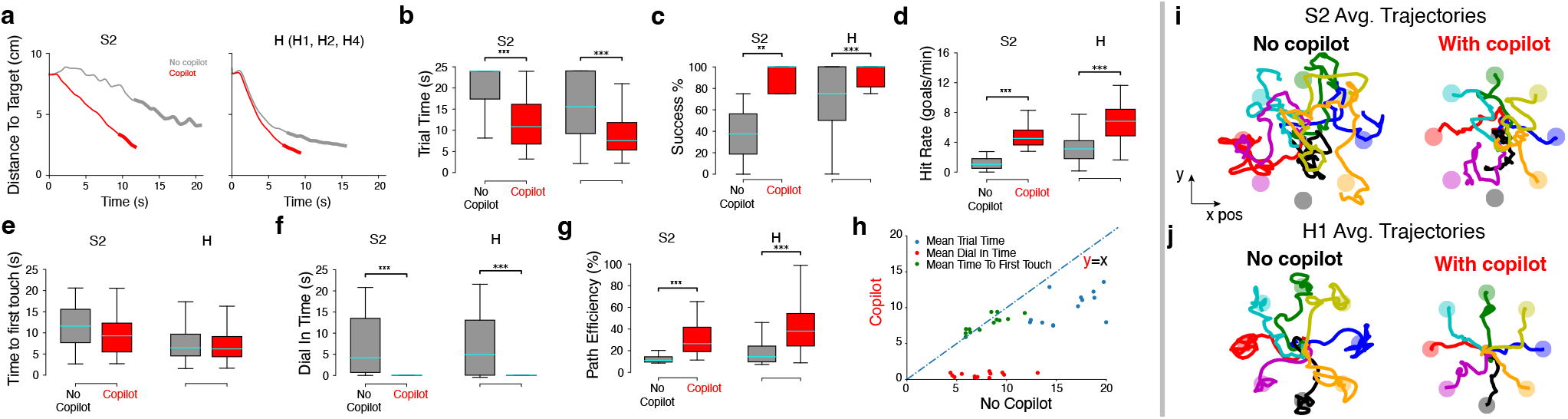
**a**, Distance-to-target plots averaged across all successful trials on the center-out 8 task for S2 and all healthy participants (H). The bolded line corresponds to the average dial-in time. **b**, Average trial time of successful radial target trials. **c**, Average success percentage of radial target trials. **d**, Average hit rate (successful acquisition of only radial target goals) per minute. Center targets, which occurred after every radial target, were not included. **e**, Average time to first touch the target for successful radial trials (applies to all following panels). **f**, Average dial-in time. **g**, Average path efficiency. **h**, A summary scatter plot of the mean trial time, dial-in time, and first touch time for all blocks, with the copilot generally improving performance. **i, j**, Average trajectories towards each radial goal with and without the copilot for S2 (**i**) and H1 (**j**). The copilot helps improve path efficiency towards the goal.

How did the copilot achieve this increased performance? First, we note the copilot was able to correctly infer the participant’s goal on every successful trial. If the copilot did not, the AI-BMI would never acquire the correct goal. Second, the copilot significantly decreased trial time. This could be achieved through two means: ballistic control (time it takes to first touch a target) or fine control (time it takes to dial-in to the target) could be improved. Although we found the median time to reach a target was less using the copilot, this difference was not statistically significant (Figure 4e). However, there was a significant reduction in dial-in time to acquire the goal, consistent with the cursor copilot aiding the acquisition of goals. In particular, whereas a traditional BMI had difficulty dialing in to a small target, the AI-BMI helped significantly reduced this dial-in time from a median of 4.74 s to a median of 0.05 s (Figure 4f). A breakdown of trial time reduction is shown in Figure 4h. Third, we found the cursor copilot also helped to make path trajectories more efficient, i.e., closer to a straight line towards the target (Figure 4g). Average trajectories for all center-out reach conditions are shown in Figure 4i, j, reflecting straighter trajectories when using the copilot compared to no copilot. Together, these results show that a cursor copilot is able to correctly infer user goals and aid their completion, increasing goal acquisition rates on a center-out 8 task by 2.3*×* to 4.3*×*.

### A robotic arm copilot enables proficient control in a sequential pick-and-place task

While some goals may be task-defined, they may also appear in random locations. In these situations, AI-BMIs can also use CV to infer goals. We used this approach to develop a robotic arm AI copilot, used in two pick-and-place tasks (Figure 5a-c). The copilot used CV to infer the location of potential goals and keep track of objects’ locations (Figure 5d). The goals were pick locations (blocks) and place locations (crosses). We inferred the locations of all blocks and crosses using a camera (Intel, RealSense D455) paired with a foundation model for open-set object detection (Grounding DINO)^30^. Our robotic arm copilot was programmed to identify all goal locations and aid control in real-time by precisely grasping a block when the robotic arm was within 2.54 cm of the block and placing any held block when the robotic arm was within 2.54 cm of a cross, or the block’s starting location. This robotic arm copilot therefore aided in executing grasp and place actions in the vicinity of CV inferred goals.

**Figure 5.**
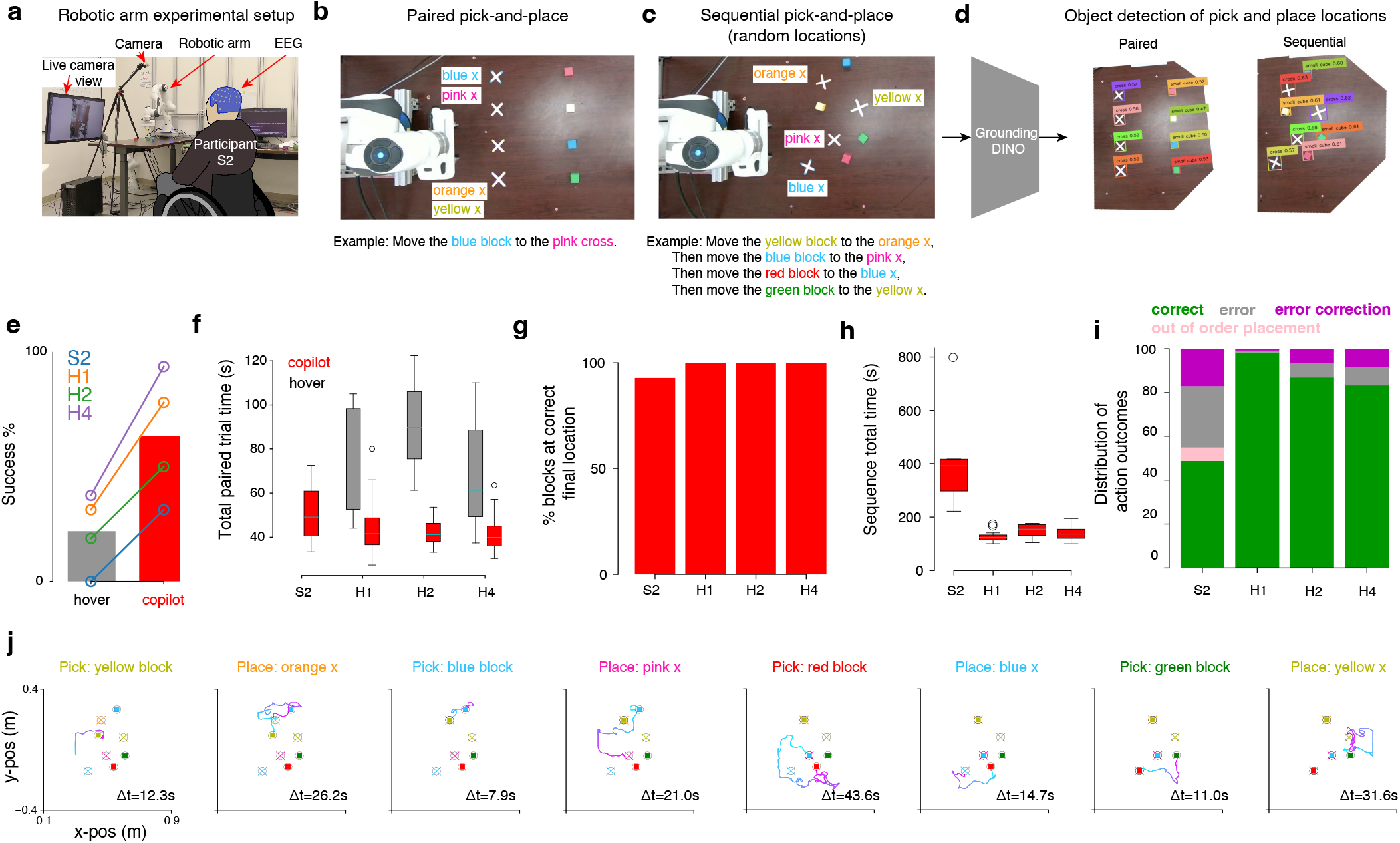
**a**, Setup of the robotic arm experiments. Each participant was seated in front of a table containing the robotic arm and a monitor showing the control orientation. **b**, Paired pick-and-place task. Participants must move a random block to a random target cross on each trial using a hover strategy or with the computer vision (CV) copilot. Inset: computer vision object identification. **c**, Sequential pick-and-place task. Each trial, the locations of target crosses were scattered, and blocks were randomly dropped onto the table. Participants were then presented with a random sequence of 4 block-target pairs, and were instructed to pick and place each block on its respective target in the order given. **d**, Computer Vision object detection using Grounding DINO for the paired (left) and sequential (right) pick-and-place tasks. **e**, Trial-level success percentage for hover and copilot conditions. **f**, Total time required to successfully pick and place blocks on their respective targets. No successful trials were recorded for S2 with the hover condition. **g**, Bar chart showing percentage of blocks that were placed on the correct target at the end of each trial of the Randomized pick and place task. **h**, Box plot showing the total time for all trials. **i**, Stacked bar chart showing the proportion of correct, out of order placements at the correct location, error, and error correction pick or place actions. **j**, Trajectories showing S2’s successful completion of the task.

We first evaluated the extent to which CV aided performance. We compared traditional BMI vs AI-BMI performance of a pick-and-place task, where 4 different colored blocks were placed in a line and 4 different colored crosses were placed in another line (Figure 5b). We called this the Robotic Arm Paired Pick-And-Place task (see Methods). On each trial, participants were asked to pick up a random block and move it to a random place location. The traditional BMI was able to give a command to pick or place via hold-time selection, while the AI-BMI relied on CV and the decoded location of the robotic arm to help execute pick and places. Trials were only successful if the participant picked up the correct block and placed it at the correct cross. Any trials where a participant picked up an incorrect block or placed it on an incorrect cross, or failed to complete a pick or place action, were counted as failures. When users were aided by the copilot, their rate of successful trials significantly increased (Figure 5e). The time to perform a trial also significantly decreased using the copilot (Figure 5f). Participate S2 was unable to perform any successful trials without the robotic CV copilot. These results therefore demonstrate that the robotic CV copilot both significantly increased success percentage and decreased trial time.

We next used this copilot to demonstrate proficient control for sequential pick-and-place tasks. We performed a task where we randomly dropped 4 blocks and 4 crosses on a table (Figure 5c). We then generated a random sequence pairing each block with a cross, and asked the participant to pick and place each block at its respective cross in order. We called this the Robotic Arm Sequence Pick-And-Place task (see Methods). We counted errors when users picked up any block that was not the next block or placed blocks at incorrect locations. These errors could be “corrected” by replacing the block at its original location, and picking up an incorrectly placed block and placing it at its assigned location, respectively. All participants successfully performed the task, placing each block at the final correct location either 93% (S2) or 100% of the time (healthy participants) (Figure 5g). The median time it took to complete a sequence (requiring 8 goals) was 392 s for S2 and 136 s for healthy participants (Figure 5h), and the majority of actions were correct actions or corrected an incorrect action (Figure 5i). Supplementary Video 1 shows S2 successfully performing this task, with trajectories plotted in Figure 5j. Together, these results demonstrate that participants are able to use a copilot to autonomously and proficiently perform a sequential pick-and-place task with a non-invasive EEG AI-BMI.

## Discussion

Our results demonstrate that AI copilots can help to significantly increase BMI control, enabling cursor and robotic arm control of a non-invasive BMI. We demonstrate a cursor and robotic arm copilot that helps users perform goal-oriented tasks. The AI-BMI increases cursor control hit rate (goals/min) by 4.3*×* for paralyzed participant S2 and enables him to perform a sequential pick-and-place robotic arm task that he was unable to do without a copilot. This approach provides a complementary axis to increase BMI performance, and as AI methods improve, this has the potential to further improve AI-BMI performance. Our improved performance was also due to stable real-time EEG decoding through an adaptive CNN-KF. We found that we only needed to decode 4 classes reliably. The four classes we chose to decode predominantly reflected different combinations of hemispheric activity near motor cortex. However, EEG performance also exhibits significant differences in performance across participants^31,32^. Further, it may be possible that in patients with ALS or other diseases, there may not be strong activation for movement-related actions due to motor neuron death. For other participants who do not exhibit decodable activity for the 4 classes we prompted, it may be necessary to identify different behavior sets that can produce 4 decodable classes. There are many candidates to try to decode 4 classes, including different limbs, wrist motor imagery^33,34^, and even cognitive imagery^35,36^. Future work should characterize how reliably 4 classes can generally be decoded based on disease condition or severity of paralysis, and increase decoding robustness. Relatedly, we note that we were unable to achieve good decoding performance with S2 on one experimental day. S2 anecdotally commented that decoding performance is inversely related with pain and paresthesia sensations, and achieved the lowest performance when pain and paresthesia were especially noticeable. As pain and paresthesia may negatively impact BMI decoding, future work should investigate how to increase EEG robustness to these factors.

We emphasize that AI copilots help reduce the required number of neurally-decoded DOFs to perform relatively high-dimensional tasks (such as the robotic arm sequential pick-and-place task). In the robotic arm task, for example, we did not require an additional click state to grasp an object; the copilot did this automatically when the robotic arm gripper was in the vicinity of the block. The use of AI copilots allows users to perform tasks beyond the complexity of a previously reported randomized grasp task^37^, where users grasped objects by holding the robotic arm’s position over each randomly placed block for 2 seconds, whose positions were explicitly made known in their control paradigm (in contrast to how our task positions were inferred via CV). For more complex tasks than these, including interacting with particular objects, training an AI copilot to handle such interactions should reduce the DOF required to handle objects. For example, a copilot controlling a robotic hand to hold a cup may fully obviate the need to decode additional DOF for finger control, something that is likely prohibitively difficult for non-invasive EEG.

Relatedly, we anticipate that future copilots may provide substantially more assistance. While our center-out copilot influenced cursor velocity at every time step, our robotic arm copilot only assisted with pick and place actions. Future copilots may also assist in executing robotic arm trajectories towards objects and precise manipulation based on the object’s dynamics. This is the goal of shared autonomy, which aims to use copilots to help increase human performance on various tasks by sharing control through task execution^38–42^. However, a limitation of these approaches is that they may potentially reduce user autonomy, leading to deleterious effects on performance and user frustration. Recently, we proposed a control sharing approach called interventional assistance that may facilitate user-copilot cooperation and increase performance on tasks^43^. Shared autonomy may enable more advanced robots, including those trained with large-scale datasets^44–47^, to act as copilots, enabling AI-BMIs to carry out even more complex movement tasks.

Recent studies have demonstrated high-performance intracortical BCI communication by decoding speech^48–50^. Our work is complementary: we focus on restoration of movement, for two reasons. First, many with paralysis (e.g., spinal cord injury below C4) do not lose speech function, but still suffer loss of movement and would be helped by motor BMIs. Second, many tasks we perform are fundamentally motor, including controlling computer cursors and robotic arms. While speech-to-movement is possible, it is less precise and responsive, as well as less discreet. For example, controlling a computer cursor through speech commands (like “move left, left, up, up, down, right”) may be prone to error with relatively crude online correction.

Future work to build better decoders may also increase performance. For example, it may be possible that large-scale EEG recordings may increase the performance of decoders, as observed in decoding language^51^ and in EMG motor decoding^52^. Because EEG is non-invasive, it is relatively easier (compared to invasive BMIs) to collect larger-scale datasets that could result in new non-invasive BMI foundation models. This approach may both reduce calibration time and increase performance.

## Acknowledgments

We thank Jason Chan, Rachel Yu, and Victoria DaSilva for participating in related pilot experiments and analyses. This work was supported by NIH DP2NS122037 and the UCLA-Amazon Science Hub.

## Author contributions statement

JYL, SL, BM, and JCK conceived of the study. JYL, SL, and BM wrote code for the real-time system, training decoders, and training copilots. JYL, SL, AM, XY, and BG conducted experiments. JYL, SL, AM, XY, BM, and BG performed analyses. JYL, BM, CK, MQ, and CX wrote code to operate the robotic arm. JYL, SL, AM, XY, and JCK generated figures and wrote the manuscript. All authors participated in manuscript review. JCK was involved with and oversaw all aspects of the work.

## Data availability

Data will be released upon publication of the manuscript.

## Code availability

Code will be released upon publication of the manuscript.

## 1. Methods

### 1.1 EEG recording

We recorded EEG signals from a 64-channel cap at a sampling rate of 1000 Hz with the eego rt amplifier (ANT Neuro, Waveguard Original, International 10/10 layout). We measured the impedance of each electrode at the start of every experimental day to ensure that none was over 200 kΩ, with a target of 20 kΩ or less. M1 and M2 electrodes were omitted from recording. EOG, Fp1, Fpz, and Fp2 electrodes were recorded but omitted from decoding because they contain strong eye movement information. We filtered the EEG data (for details, see 1.5), which was then sent as inputs to our real-time system, named Real-time Synchronous Python (RASPy).

### 1.2 Real-time experimental setup

Our open loop task operated at 100 Hz (10 ms “ticks”) and our closed-loop tasks operated at 20 Hz (50 ms ticks). These tasks are described below. In tasks with real-time decoding, we decoded the most recent 1000 available samples (1 second) of EEG data. Cursor control tasks were displayed on a 20-inch monitor (48 cm *×* 27 cm) with a refresh rate of 60 Hz. Each participant was seated with their eyes positioned approximately 55-76 cm from the screen. We recorded eye gaze position at 60 Hz (Tobii Pro Nano). The eye tracker was calibrated at the start of each day and whenever large changes in seating position occurred between sessions. Robotic arm control tasks were performed using the Panda arm (Franka Emika) running at 1000 Hz.

### 1.3 Tasks

Participants performed up to five tasks. Two tasks, “Open Loop Text” and “Decorrelated Closed Loop” were used to train decoders. One task, “Center-Out 8 Target Selection,” was used to quantify decoder performance for cursor control. Two tasks, “Robotic Arm Paired Pick-And-Place” and “Robotic Arm Sequence Pick-And-Place” were used to quantify decoder performance for robotic arm control. Participants were asked to sit up straight and rest their hands in a comfortable position for all tasks.

#### Open Loop Text (referred to as the “open loop” task)

We presented text corresponding to four actions. We randomized the presentation of the actions in groups of 4. We refer to the presentation of one action as a “trial.” Each trial was 20 seconds long. The participant was instructed to repetitively perform an action corresponding to the presented text. There was a 2 second inter-trial interval. During the inter-trial interval, a red box was displayed and the participant was asked to relax and not to perform any action. The presented texts for the healthy participants were: Left hand, Right hand, Both hands, or Feet. Healthy participants overtly moved the limb corresponding to the presented text (for more details on why we used overt movements in healthy participants, please see Methods 1.4). The presented texts for the spinal cord injury (SCI) participant were: Left leg, Right leg, Both legs, or Still. The SCI participant was asked to attempt to move the paralyzed limbs or to remain still during the respective prompts. The SCI participant displayed no observable limb movements.

#### Decorrelated Closed Loop (referred to as the “decorrelated” task)

The goal of this task is to decorrelate eye movements from motor actions. Removing the effect of eye movements is critical, since signals reflecting eye movements are present in EEG. The goal of our BMI was to decode motor activity, not eye movements. We therefore designed this task so that trained decoders could accurately decode motor activity but not eye movements. This task utilized a decoder trained from the open loop task that controlled the position of a cursor. We provided the user with real-time feedback of the cursor position.

At the start of a decorrelated trial, a cursor was displayed at the center of the screen. A text prompt (from the same set used in the open loop task) appeared inside a rectangular box at one of eight target locations (from 0^*°*^ to 315^*°*^ in 45^*°*^ increments, centered at a radius of 9.34 cm and width of 4.67 cm). The text prompt and its target location were uncorrelated. For example, ‘Left Leg’ could appear at any of the eight locations for that trial. The output from the decoder trained on the open loop data was used for the cursor’s movement in the decorrelated session. The decoder was continuously adapted (see Methods 1.7 for more details). The cursor’s movement was restricted to move along the straight line segment connecting the center of the screen to the outer edge of the target. To do this, we decoded 2D velocity and rotated the velocities to be congruent so that the correct action moved the cursor towards the correct target. For example, if ‘Left Leg’ was at the up right target (45^*°*^, Figure 2h) then any left cursor movements moved the cursor up right. The 2D velocities were subsequently projected onto the straight line segment connecting the center of the screen to the outer edge of the target. Intuitively, the cursor moved toward the target if the decoded action was the same as the prompted action. The cursor moved away from the target if the decoded action was opposite.

Although a target was presented in the decorrelated closed loop task, the target was not selectable, and remained onscreen for 7 seconds, with cursor movement occurring only in the last 5 seconds. We chose to not make targets selectable to acquire an equal number of samples per class in the training data. If targets were selectable, then actions that were more accurately decoded would be underrepresented in the decorrelated task data. If the decoder continued to decode the correct action, the cursor would hover within the target acceptance window for the duration of the trial.

All decoders used for closed-loop control were trained using data from the decorrelated task to minimize the influence of eye position on decoding. A performant decoder must therefore *ignore* eye artifact, since eye movement signals more frequently correspond to classes other than the correct class, and would be detrimental to the decoder.

#### Center-out 8 Target Selection (referred to as the “center-out 8” task)

During the center-out 8 task, the user controls a cursor to acquire 8 targets equally spaced on the circumference of a circle. The cursor was displayed at the center of the screen in grey color. The position of the cursor was bounded to a square with *x*-position between [*−*1, 1] units and *y*-position between [*−*1, 1] units (total side length: 2 units). 1 unit corresponded to 11.675 cm. The target appeared in green color at one of the 8 locations spaced 0.7 units from the center at 45^*°*^ intervals. The target diameter varied from 0.6 units (7 cm) to 0.2 units (2.3 cm). To acquire a target successfully, the decoded cursor had to be held contiguously over the target for 500 ms. The trial timed out if the participant was not able to acquire the target within 24 seconds. After each center-out target, the next prompted target was at the center of the screen. If the participant wasn’t able to acquire the center target, the cursor was reset to the center and the target was placed at one of the center-out locations. If a trial ended with target acquisition, it was counted as a success. If the trial ended after timeout, it was counted as a failure. Statistics for the center-out 8 task are only computed on the 8 center-out trials to be agnostic to target location and are averaged over each set of 8 trials.

#### Robotic Arm Paired Pick-And-Place (referred to as the “Robotic paired” task)

We setup four place locations (crosses, spaced approximately 13 cm apart along a line) and four pick locations (2.9 *×* 2.9 *×* 2.9 cm^3^ blocks, also spaced approximately 13 cm apart, and approximately 28 cm away from each cross). We then generated a trial where a randomly chosen block was paired with a randomly chosen cross. To successfully complete a trial, the participant had to control the robotic arm to (1) move to the block, (2) grasp the block, (3) move the block to the correct cross, and (4) place the block on the cross. The trial was otherwise failed. The participant had 40 seconds to grasp the block before the trial was failed. If the participant grasped the block, they would then have 40 seconds to place the block before the trial was failed. Trials were performed in sets of 8. For the first 4 trials of each set, participants performed all 4 trials with (or without) an AI copilot. For the last 4 trials of each set, participants received the same prompt as the first 4 trials of the set, and performed the trials without (or with) an AI copilot. Whether each set began with or without an AI copilot was randomly determined (by a coin flip with *p* = 0.5). Participants were informed whether they performed the task with or without an AI copilot.

The user controlled the 2D endpoint of the robotic arm above the blocks and crosses, with altitude as a function of distance to the closest block/cross. Users were shown a live camera view from an overhead camera of the workspace (Figure 5a). Users were also given the option to rotate this top-down view of the workspace; S2 chose *up* and H1, H2, and H4 chose *left* to correspond to the direction pointing away from the base of the robot. The maximum 2D speed of the robotic arm was set to 5 cm/s. When the computer vision (CV) AI copilot was used, it would grasp (place) a block whenever it was within a 2.54 cm radius of any block (cross). When CV was not used, the robotic arm would attempt a grasp (or place) when it stayed within a radius of 6 cm for 4 seconds. When the grasp was attempted over a region that did not have a block, the arm lowered, closed and opened its gripper, then returned to an altitude above the blocks. When the arm collided with a block while lowering, it stopped attempting the grasp (or place) and returned to an altitude above the blocks. When a place was attempted over a region that did not have a cross, the arm proceeded to place the block at that location. The time for a grasp or place attempt was not counted towards the 40 second timeout time for either grasping or placing a block; only time while the participant freely moved the arm counted toward the timeout time.

#### Robotic Arm Sequence Pick-And-Place (referred to as the “Robotic sequence” task)

During the pick-and-place task, the user controlled a robotic arm to pick and place 4 colored blocks (“pick” locations) onto 4 crosses (“place” locations). The 4 place locations were randomly selected. The 4 blocks were then randomly dropped on the table, and only moved if a block was out-of-bounds of the workspace. A random sequence was generated, assigning each of the 4 blocks uniquely to the 4 place locations in a particular order. The subject had to control the robotic arm endpoint to place each block at each cross in the specified correct order.

### 1.4 Experimental Protocol

#### Participant selection

All experiments were approved by the UCLA IRB. Three healthy participants and one paralyzed participant with spinal cord injury (SCI) (paraplegic, T5 complete) participated in the full duration of this study. As stated in the Results and Discussion, controlling the CNN-KF used 4 decodable classes to control right, left, up, and down 2D cursor movements. Further, participants demonstrate varying levels of EEG modulation for particular actions^32^ that can be affected by several factors, including mind-body awareness^31^. While any 4 decodable classes could be used to control our BMI, we only tested the set of prompted behaviors described in the task description of Open Loop Text (for healthy: left hand, right hand, both hands, legs; for SCI: left leg, right leg, both legs, still). If a participant did not achieve high decoding accuracy on these 4 actions from the open loop task, we did not perform subsequent experiments to identify a behavior set that generated 4 decodable states, as this search was open-ended. Rather, we decided to only perform experiments on participants whose activity for the prompted action set could be decoded by a CNN-KF.

Two additional SCI paraplegic participants were evaluated for a single day, but did not achieve high decoding performance for attempted leg movements. This does not mean these participants could not control a BMI, but only that they did not exhibit substantial EEG modulation for the prompted behaviors. To be clear, we did not perform further experiments to identify a behavior set that could produce 4 decodable classes, since determining these behaviors requires an open-ended search over many experiments, a tangential question representing future work to make EEG BMIs more robust (see Discussion). Two additional participants (one SCI, the other healthy) conducted pilot experiments, achieving 2D cursor control, but did not complete experiments for personal reasons unrelated to the study.

#### Training data collection

Each participant performed the open-loop task for approximately 20 minutes. EEG activity corresponding to each of the four possible motor actions was used to train a CNN-KF (see 1.6, 1.7). There were two important differences between the open loop and the closed loop cursor control sessions. First, there was real-time feedback on the decoder’s output in the form of the cursor movement. Second, user’s eyes followed the cursor during closed loop cursor control, resulting in eye movement correlated to the task at hand. To ensure that the KF was not adapting to eye movement EEG artifact during the cursor control session, we included a decorrelated closed loop session so that the EEGNet CNN learned to ignore the eye movements.

Analysis showed that each user’s eye movements during the decorrelated task were minimally correlated with cursor and target information (Extended Data Figures 3 and 4), thereby minimizing the effective contribution of eye movements to decoding. A new EEGNet was trained using the data from the decorrelated session. This trained EEGNet was used in the following center-out-8 closed-loop sessions. To control the movement of the cursor, the SCI participant was asked to do the action tasks: ‘Left leg’ for left, ‘Right leg’ for right, ‘Both legs’ for up, and ‘Still’ for down. The healthy participants moved ‘Left hand’, ‘Right hand’, ‘Both hands’, and ‘Legs’ to move the cursor to the left, right, up, and down respectively.

Although our SCI participant did not have any overt movement of his paralyzed limbs during BMI control, we instructed healthy participants to overtly move their hands, in contrast to imagined movements. Overt movements have been previously used in several monkey BMI models^11,53^ that were successfully translated to paralyzed human participants^2,3^. Our choice of overt versus imagined movements for healthy participants centers around different considerations in using able-bodied participants to model paralyzed participants, something also considered in intracortical BMI animal models^54^. While overt movements are limited in that there is proprioceptive feedback^55^, a concern of imagined movements is that they correspond to significantly constrained motor cortical activity. Imagined movements do not produce movement, meaning they are constrained to a lower variance “output null space” of motor cortical neural population activity^56^. A significant proportion of motor cortex variance is related to neural activity that generates movements^57^. In attempted movements, paralyzed participants can generate activity that would have produced movement but does not ultimately result in movement due to their injury or disease (e.g., spinal cord injury, meaning movement generating neural signals do not reach the muscles). To not significantly constrain motor cortex activation, we decided to record “output potent” neural activity that produces movement, a signal still present in paralyzed participants. We therefore decoded EEG activity corresponding to overt movements (healthy participants) or attempted movements with no overt behavior (paralyzed participant). Please note that all of our BMI experiments worked with S2, who displayed no overt movement of his paralyzed limbs during BMI control.

### 1.5 Data preprocessing

We filtered EEG data using a 4th order Butterworth low-pass filter (cutoff 40 Hz) cascaded with a 5th order Butterworth high-pass filter (cutoff 4 Hz) and notch filters at [60 Hz, 60 Hz, 120 Hz, 180 Hz] with quality factors of [10, 4, 5, 2] respectively. Following this, we applied a band-pass 5th order Butterworth filter to each of the remaining channels, allowing frequencies between 4 *−* 40 Hz to pass through. This range contains the typical brain oscillations of interest, such as theta, alpha, and beta waves, while attenuating slow drifts and high-frequency noise. We then employed a next nearest neighbors Laplacian spatial filter over decoded electrodes. Finally, we normalized the data for each channel to have unit root-mean-square (RMS) for faster convergence in the subsequent neural network. This normalization aims to increase robustness to signal amplitude variance that occurs across recording sessions.

#### 1.6 EEGNet (CNN) training

We used a convolutional neural network, EEGNet^21^, to classify EEG signals. First, the preprocessed EEG data was fit with 8 temporal filters of size (1, 51), outputting 8 feature maps containing the EEG signal at different band-pass frequencies. The filter size of 51 was chosen to be just over half of the downsampled EEG frequency of 100 Hz (`scipy.signal.resample`). This convolution layer was followed by a batch-norm layer. We then used 2 spatial filters of size (*C*, 1) where *C* = 58 is the number of channels in the EEG signals. This spatial layer was followed by a batch-norm, average pool, and dropout layer. The filter size for the average pool layer was set to (1, 3) to reduce the sampling rate of the signal to around 33 Hz. The output from the average pool layer was fed into a separable convolution layer – depthwise convolution (size (1, 16)) followed by a (1, 1) pointwise convolution. This separable convolution layer was followed by a batch-norm, average pool (1, 16), and dropout layer.

Custom Python code was used to implement and train the EEGNet. We split the data into 5 folds at the trial level, and performed 5-fold cross-validation over these folds. The learning rate was set to 10^*−*3^ at the start, and reduced the learning rate by a factor of 0.1 if the accuracy on the held-out validation fold did not improve for 10 consecutive epochs. We also halted training if the minimum validation loss did not improve for 10 consecutive epochs. From the 5-fold cross-validation, we chose the model for online control based on the validation confusion matrix and overall accuracy.

Additionally, for compatibility with the Kalman Filter so that error distributions more closely resembled the Gaussian distribution, the EEGNet models were trained with a scaled mean-squared error (MSE) loss with targets corresponding to one-hot vectors of each class. The MSE loss was scaled by a factor of 100 to improve training stability. Accuracy for models trained with the one-hot mean-squared error loss was defined by labeling the prediction of each data point as the argmax over the four classes. This method is equivalent to a Linear Discriminant Analysis classifier with fixed mean and covariance and with trained feature maps. Data used to train the EEGNet was also used to train a seed Kalman Filter for later closed-loop sessions.

#### 1.7 Kalman filtering

The discrete-time linear Kalman filter (KF) with no inputs assumes a state-observation linear dynamical system defined as:

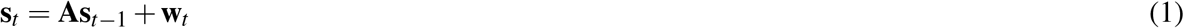

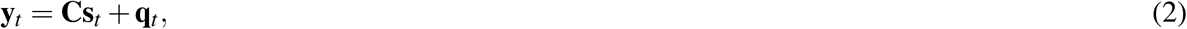

In this instance, **s**_*t*_ is the cursor (or robotic arm) kinematic state vector at time step *t* representing position and velocity, **y**_*t*_ is the observation vector (EEGNet hidden state), **A** and **C** are the state transition and observation matrices, respectively, and **w**_*k*_ and **q**_*k*_ are state and observation noise. The KF decodes according to

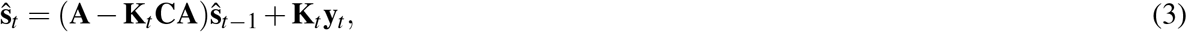

where **K**_*t*_ is the Kalman gain at time *t*. The linear KF is minimum MSE optimal when noise **w**_*t*_ and **q**_*t*_ are Gaussian and independent of time.

To adapt this formulation to four-class decoding, we separate velocity into 2 components per axis: *v*^+*x*^, *v*^*−x*^, *v*^+*y*^, and *v*^*−y*^. A positive *v*^+*x*^ corresponds to a positive *x* axis velocity while a positive *v*^*−x*^ corresponds to a negative *x* axis velocity. The kinematic state vector is then defined as

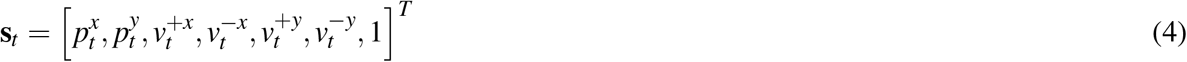

for *x* position *p*^*x*^ and *y* position *p*^*y*^ (a.u.). At each time step, we additionally clip the *L*_1_ norm of velocity components 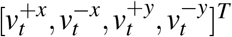 to be at most 1.

We set **A** and **W** to

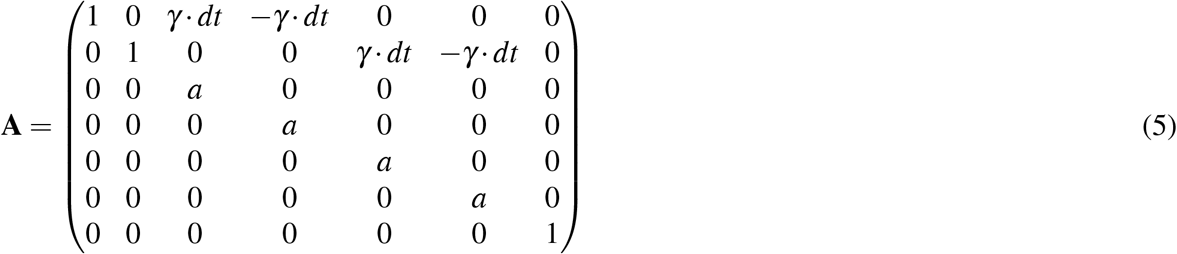

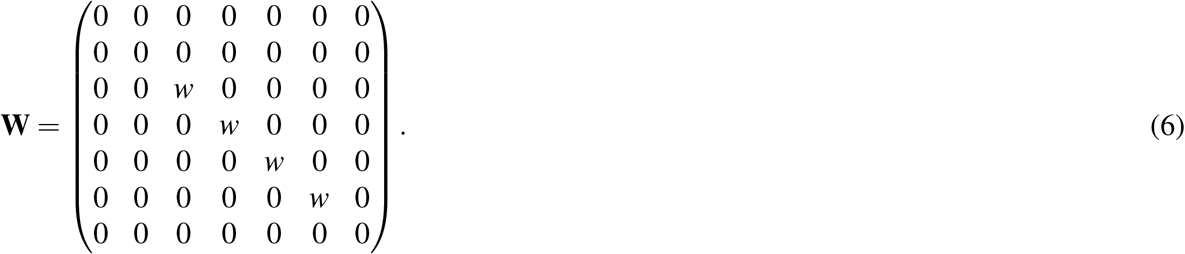

The gain *γ* was set to 0.5, *dt* was the tick length 50 ms, *a* was set to (0.825^1*/*0.2^)^(*dt*)^, consistent with Silversmith *et al*.^14^, and *w* was set to the MSE loss averaged over all dimensions from training EEGNet. Additionally, for stability during decoding, we normalized the L1 norm of velocity components to have a maximum value of 1 prior to performing the state update. This effectively sets a maximum velocity of 5.84 cm/s (0.5 a.u./s).

### 1.8 Closed loop decoder adaptation

The hidden state of the EEGNet trained from the decorrelated task was used for the observations of a linear KF, whose state represented the position and velocity of the displayed cursor. We performed closed-loop decoder adaptation to adapt the Kalman Filter through the course of each session^58^. While internal statistics were updated at each tick, online decoder parameters were only updated after each trial of an adaptation block or after the end of the first evaluation block. Except for some earlier sessions (H2 Day 1-5, S2 Day 1), the position information was removed from the inferred state prior to adaptation, so that the Kalman Filter was effectively a Velocity Kalman Filter^11,15^. Later sessions for S2 were readapted to past data with all position information removed from the inferred state. For all sessions, the rows of the observation matrix **C** corresponding to position were set to zero. As with Gilja *et al*.^11^, the rows and columns of the a posteriori estimate covariance matrix were set to zero at every iteration.

Retraining or CLDA for continuous BMIs significantly benefits from a ReFIT-KF innovation that augments the trianing data by defining the velocity component of the inferred state as a vector pointing from cursor to target. However, in our context, when the user is commanding one of four actions rather than continuously attempting to move their arm, this may not be optimal. For example, when trying to reach the up right target (45^*°*^) from the origin, both an up and right command are equally correct. Instead of a single state at each timestep, the inferred state is represented by a probability distribution over up to 2 plausible states. For example, when the cursor is at coordinates (0, 0) and the target is at coordinates (0.15, 0.35), we assign probability of 0.3 to the state [0, 0, 1, 0, 0, 0, 1]^*T*^ (right) and probability 0.7 to state [0, 0, 0, 0, 1, 0, 1]^*T*^ (up). Formally, and dropping the time step subscript for convenience, the inferred state was defined by

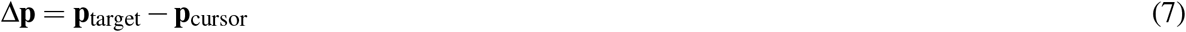

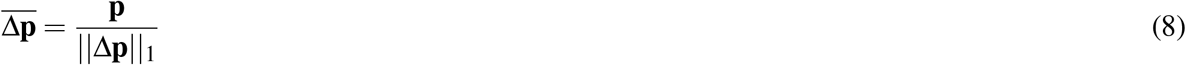

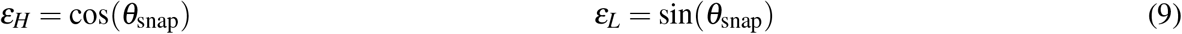

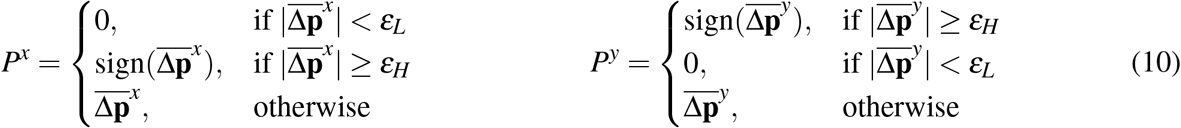

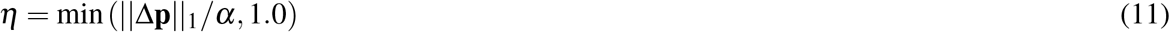

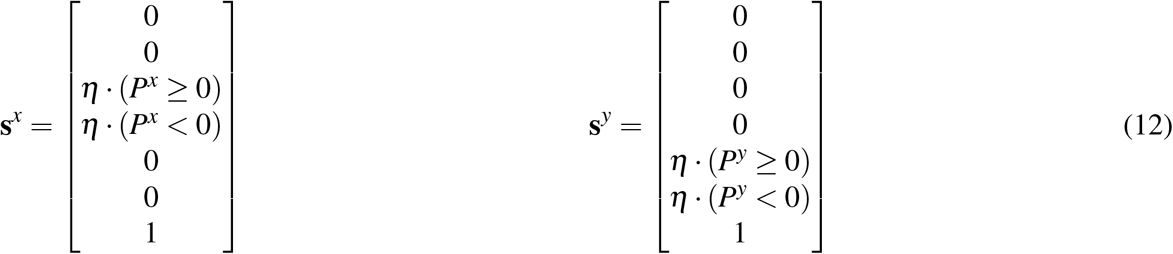

where **p**_target_ and **p**_cursor_ are the (*x, y*) positions of target and cursor, *θ*_snap_ = 22.5^*°*^ is the angle under which vectors are snapped to the nearest direction, **s**^*x*^ and **s**^*y*^ are the states corresponding to velocities on the x and y axis, respectively, and *η* acts as a scale factor parameterized by *α* = 0.2 a.u. The *x*-axis state **s**^*x*^ was represented with probability |*P*^*x*^|, and **s**^*y*^ was represented with probability |*P*^*y*^|. During the decorrelated closed loop session, inferred states were set as [0, 0, 𝕀(right), 𝕀(left), 𝕀(up), 𝕀(down), 1]^*T*^, e.g. [0, 0, 0, 0, 0, 1, 1]^*T*^ for a prompt corresponding to the down action. Future work to optimize the inferred state could further improve performance.

On some days, closed-loop performance of a previous day’s decoder dropped significantly during a center-out-8 session. On those days, we collected data from a new decorrelated training session to train a new EEGNet. We then adapted the KF resulting from the new EEGNet to a center-out-8 dataset that had already been collected (S2 Day 4 decoder adapted to Day 3 data). This reduced the amount of recalibration time needed when a new EEGNet was trained. The authors note that while past data from center-out and other 2D tasks could be used to train an EEGNet with more data, they did not do so in this work in order to minimize the effect of eye movements on decoding.

### 1.9 Cursor copilot

#### 1.9.1 Overview

The cursor copilot aids the user’s cursor control by directly influencing cursor movement with its own copilot output velocity 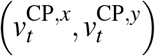 which is added to the cursor velocity derived from CNN-KF 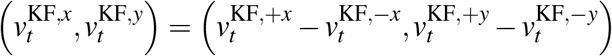.

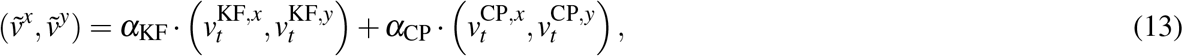

with *α*_KF_ = 0.6 and *α*_CP_ = 0.3.

The copilot used a Long Short-Term Memory (LSTM) recurrent neural network to learn representations over historical inputs. The input to the copilot network was the cursor’s current position and velocity. The goal information (i.e., the prompted target) was not known to the copilot. Rather, the copilot had to infer the user’s goal (one out of the 9 candidate center-out 8 targets) from the user’s past movements. Given the dynamic task information, the copilot therefore implicitly inferred the user’s intended goal and influenced the cursor’s velocity to move more efficiently toward the goal.

#### 1.9.2 Input and output

For our experiment, we made a design choice of using a policy network that input the CNN-KF velocity and the cursor’s trajectory to output copilot velocity 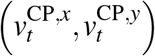. The LSTM component of the policy network has recurrence, enabling it to model reperesentations of the cursor trajectory. The copilot thus only observed the cursor positions and the current KF velocity, 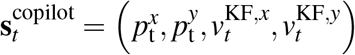 at every timestep. At the beginning of each trial, the LSTM hidden state was reset to zeros. These values updated as the cursor moved. Overall, the LSTM network mapped an observation 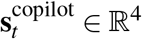 to an action in **a**_*t*_*∈*ℝ^9^. This action influenced the movement of the cursor by using a charge-based system with electric charges on all 9 target positions. To define this charge-based system, we first define a matrix **p**_targets_*∈*ℝ^2*×*9^ defining the (*x, y*) positions of all 9 targets:

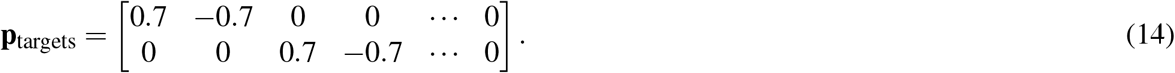

As described in the main manuscript, the center-out 8 task has 9 defined targets that are fixed. Our design incorporates this task specific information, however, in general computer usage, this copilot can use candidate target locations **p**_targets_ inferred via computer vision. We let 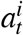denote the *i*th element of the and note that every element of **a**_*t*_, output by the LSTM. Each element 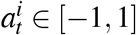. We then define the “charge” assigned to target *i* at time *t* be

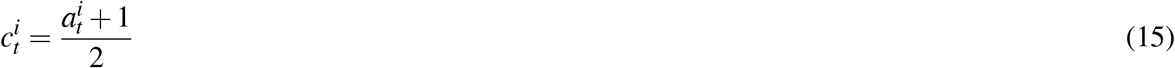

meaning each charge 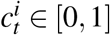. We then use these charges to define a “force” that each target exerts on the cursor, mimicking Coulomb’s law. We let hyperparameter *q* denote the charge of the cursor, and compute the force exerted by each of the 9 targets via:

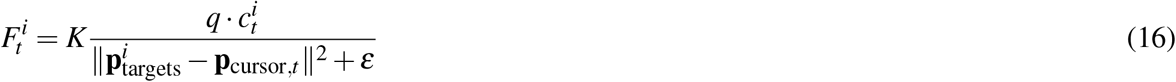

where 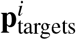 is the *i*th column of **p**_targets_, **p**_cursor,*t*_ is the current cursor position at time *t*, hyperparameter

*K* is a fixed constant, and hyperparameter *ε* avoids division by zero. The value of all hyperparameters are given in Extended Data Table 3. We then compute the charge exerted by the *i*th target on the cursor at time *t* as

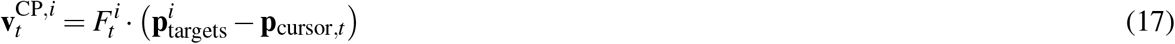

which is a vector *∈* ℝ^2^. The total influence of all targets on the cursor is

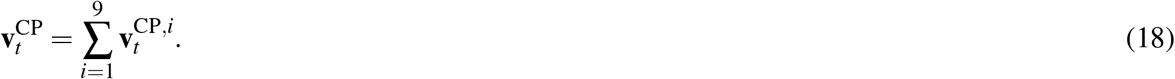

This sum was the copilot output velocity, 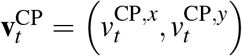 that influenced the cursor towards the inferred goal.

#### 1.9.3 Cursor task rewards

The policy network was trained via Proximal Policy Optimization (PPO)^29^. The reward signal at each timestep, *r*_*t*_, was defined by the proximity of the cursor to the target, where

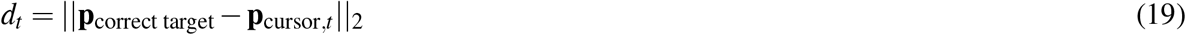

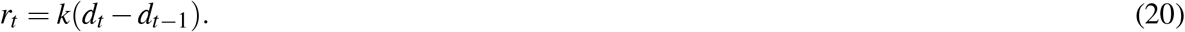

The hyperparameter *k* was chosen so that the reward, *r*_*t*_, in each trial was between *−*1 and 1. Please note that the cumulative reward is a state function. Future rewards were discounted at *γ* = 0.989 to encourage the copilot to acquire the target in the fewest number of steps. The copilot value network (*V* (**s**)) estimated the expected rewards-to-go from state **s**.

#### 1.9.4 Network architecture and training hyperparameters

The value, *V* (**s**), and policy, *π*(**s**), networks were composed of a two-layer multi-layer perceptron (MLP) with 64 hidden units, followed by an LSTM with a 256-dimensional hidden state. The copilot was trained using PPO with a fixed learning rate of 3 *×* 10^*−*4^ with a rollout buffer size of 2048 and a batch size of 64. Copilots were trained for a total of 1, 200, 000 steps.

#### 1.9.5 Training

Training the copilot involves generating decoded velocities, 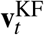, from the CNN-KF. However, optimizing the copilot with online human-in-the-loop experiments is very expensive in time. We therefore modeled CNN-KF decoded velocities to accelerate copilot training. We call the model that generates CNN-KF decoded velocities the “surrogate human control policy” because it models decoded velocities from human BMI users. The surrogate human control policy, denoted 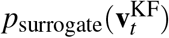, approximates the distribution of human cursor control velocities, 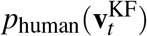, when performing the center-out 8 task. We designed the surrogate policy to mimic four features of the human policy:

1. The human attempts to move the cursor in the correct direction. To model this, we defined an intended velocity,

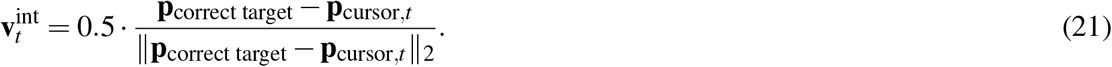
2. Due to the slow dynamics of EEG activity, changing cursor directions only occurred after a modest delay. We therefore modeled a switch in decoded direction at time *t* as occurring over an interval between [*t, t* + delay] and linearly interpolated velocities between 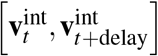. The random delay was drawn from a uniform distribution, *U* [300*ms*, 600*ms*].
3. When a new direction is decoded, the CNN-KF decoded velocities can be erroneous, which we modeled via angular noise in the decoded velocity. We modeled this as:

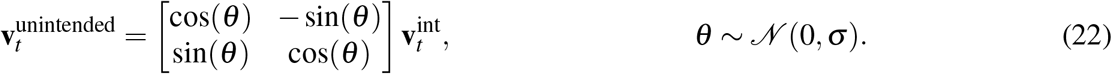

over the switch direction interval [*t, t* + delay]. In our simulations, *σ* = *π/*3.
4. Even when CNN-KF decoded velocities go in the correct direction, there is stochasticity in cursor movements. We modeled this via independent additive Gaussian noise. This produced the final surrogate simulated velocity,

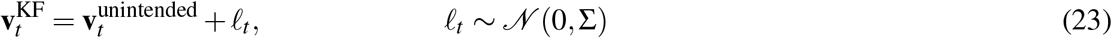

In our simulations, Σ = 0.03**I**.

With this model of 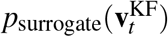, we trained the copilot without a human-in-the-loop. In copilot training, we also increased the hold time to 2 s to encourage the copilot to hold the cursor close to the target even after acquiring it. All hyperparameters are reported in Extended Data Table 3.

- **Center-out 8 performance metrics**

We report how each performance metric for center-out 8 was computed.

- **Trial Time (s)**: the time from trial initiation to trial completion.
- **Success** %: the percentage of correctly acquired radial targets.
- **Hit rate (goals per minute)**: The number of successfully acquired radial targets (goals) per minute. No part of the trial was excluded.
- **Time to first touch (s)**: the time it takes the cursor to first touch the prompted radial target.
- **Dial-in time (s)**: the time between when the cursor first acquires and last acquires the prompted radial target^11^. If the BCI user successfully holds the target when first acquiring it (i.e., stays on it for 500 contiguous ms after first touching it), the dial-in time is zero. If the cursor *moves off* the target after first touching it, then the dial-in time will be greater than zero because the time when it last acquires the target (prior to the 500 ms contiguous hold time) is different from when it first acquired the target. To be clear, dial-in time does not include the 500 ms hold time; it only includes the time spent “dialing in” on the target to acquire it.
- **Path efficiency** (%): the straight line distance between the targets divided by the total distance traveled by the cursor, multiplied by 100%. Its maximum value, 100%, occurs when the cursor takes the shortest path between targets.
- **Fitts’ Information Transfer Rate (ITR, bits per second)**: Fitts’ ITR^59^ is defined as

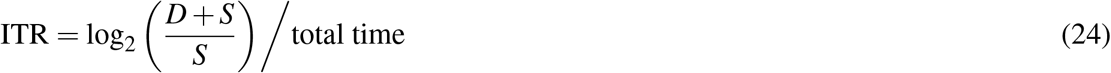

### 1.11 Spatial saliency maps

To visualize the contribution of each EEG channel’s activity toward decoding, we computed saliency maps over channels. Saliency maps represent the importance of input channels for driving the cursor in one of the four cardinal directions (right, left, up, and down). Spatial saliency maps for each direction were calculated as

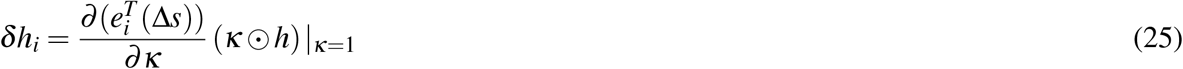

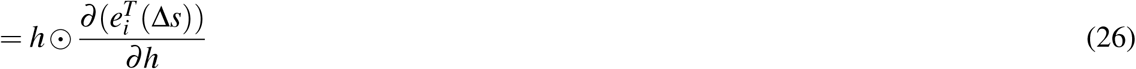

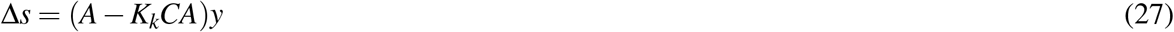

where *h* is an input or intermediate layer activation of the decoder, *e*_*i*_ is the unit vector for index *i, κ* is a unit scaling factor to collect gradients, and is ⊙ the broadcasted Hadamard product operator. We averaged *δh*_*i*_ over the time dimension to produce saliency maps for preprocessed inputs and for inputs to the EEGNet depthwise layer. Saliency maps were then projected using MNE’s `plot_topomap` function^60^.

### 1.12 Robotic arm and computer vision copilot

We interfaced with the Franka Emika Panda 3 robotic arm using the frankalib and liborl C++ libraries with custom C++ and Python code, mounted above a table. A camera (Intel RealSense D455, 1280×800px, 30Hz, RGB) was positioned with a tripod such that it viewed the working space of the robotic arm. We calibrated the computer vision system using 4 marked locations on the table, and masked the image corresponding to the working space. A separate process used GroundingDINO^30^ to find the pixel locations corresponding to the “small cube” and “cross” text prompts. This typically took 350-400ms on an RTX A4000 and can be extended to arbitrary objects. For safety, the robot’s (*x, y*) position was limited to be between the polygon defined by vertices (0.3, *−* 0.31)m, (0.6, *−* 0.31)m, (0.75, *−* 0.15)m, (0.75, *−* 0.15)m, (0.6, 0.31)m, and (0.3, 0.31)m. Participants performed two tasks: a fixed location pick-and-place control task with or without computer vision and a 4-block-target sequence pick-and-place task. In each task, participants placed 4 colored cubes (green, blue, yellow, red, 2.9 cm width) on colored cross markers (yellow, orange, pink, blue, 5.4 cm width). We use the term “goal” to refer to both pick locations (blocks) and place locations (crosses).

In the control task, for each trial, participants received a prompt of which color cube to place at which color target cross location. The user had to perform the task for each of two conditions: *hover*, in which remaining within a horizontal radius of 6 cm of the robot’s average (*x, y*) position over 4 seconds initiated a pick/place at the average location, and *computer vision copilot*, in which entering within a 2.54 cm radius of a goal initiated a pick/place action at the goal’s location. In the *hover* condition, the average location was continuously updated, allowing for minor finetuning of the final pick/place location. A 5-second cooldown for initiating a new pick/place action was enforced after the end of each pick/place action. 4 blocks and 4 target crosses were placed at [(0.675, *−* 0.179), (0.674, *−* 0.058), (0.672, *−* 0.075), (0.674, 0.207)]m and [(0.387, *−* 0.174), (0.388, *−* 0.052), (0.392, 0.087), (0.395, 0.210)]m and were reset in between trials (12-14cm between adjacent blocks/targets, and 28cm between each block and the closest target). The robotic arm’s end effector was reset to (*−*0.3, 0.021, 0.081)m before each trial. The z-position of the robotic arm was approximately (−0.03 + 0.11 *·* min(5 *·* (*d −* 0.0254), 1))m, with the distance to closest eligible block/target *d* in meters. Each trial began when a button to allow movement was pressed and ended when the user placed a block at any location or when 40 seconds of moving time (time not dedicated to a pick/place action) accumulated. We interleaved pairs of sets of 4 trials for *hover* and *computer vision copilot* conditions, with the order of each pair randomly generated, and with the same 4 prompts for each condition. Users were informed of which condition was active so that they could adjust their control strategy accordingly.

For each trial of the sequences task, after resetting the robotic arm’s position, 4 targets (colored crosses) were randomly positioned within the working space, then 4 colored blocks were randomly dropped onto the table. If any of the blocks was outside of the reachable area of the arm, or was too close to any other block or target (approximately 5 cm), the block was moved to a nearby valid location. The computer vision system detected each block and target’s position on the table. These positions were set once movement began and were updated according to the robotic arm’s state, i.e. keeping track of the current locations of blocks and if targets were occupied. The system did not allow placement of new blocks onto already occupied targets, but did allow blocks to be placed at the location from which they were picked up. We then generated a list of block-target pairs, and instructed the user to place each block at its respective target in that order. If the user picked up a block out of order, they were instructed to replace the block at its original location. If this was subjectively too difficult or not possible (due to the occupation of place locations), they were instead instructed to place the out-of-order block at its respective target location.

Whenever the (*x, y*) position of the robotic arm was within 2.54 cm of the detected center of each block or target location, the robotic arm automatically picked up or placed the block on the table. Trials ended when all 4 blocks were placed, when 40 s of moving time elapsed and the trial timed out, or when users declined to finish the trial. On some trials, the gripper dropped the block or failed to place the block due to improper calibration. These trials were excluded from analysis.

On earlier trials for S2 sequence task (4/7 trials), the maximum speed was only 4 cm/s, and the robotic arm only picked/placed the block when the end effector was within 2.54 cm of the block/target and the block/target had the highest score = (*r*_*target*_*− r*_*robot*_) *·* ∠(*v, r*_*target*_ *− r*_*robot*_), where ∠(*v, r*_*target*_ *− r*_*robot*_) is the angle between the 2D KF velocity *v* and 2D displacement *r*_*target*_ *− r*_*robot*_. This resulted in 2 instances on 2 trials where the robotic arm failed to place a block on a target which it entered the radius of (time difference 18.4 s, 10.7 s) and 1 instance on the first of the aforementioned trials where the robotic arm failed to pick up the correct next block in the sequence and instead picked up an incorrect block (time difference 5.3 s). Since these instances decreased performance, we did not modify statistics for these trials.

## References

1. Hochberg, L. R. et al. Neuronal ensemble control of prosthetic devices by a human with tetraplegia. Nature 442, 164–171 (2006).

2. Gilja, V. et al. Clinical translation of a high-performance neural prosthesis. Nature Medicine 21, 1142–1145 (2015).

3. Pandarinath, C. et al. High performance communication by people with paralysis using an intracortical brain-computer interface. eLife 6, e18554 (2017).

4. Hochberg, L. R. et al. Reach and grasp by people with tetraplegia using a neurally controlled robotic arm. Nature 485, 372–375 (2012).

5. Collinger, J. L. et al. High-performance neuroprosthetic control by an individual with tetraplegia. The Lancet 381, 557–564 (2013).

6. Wodlinger, B. et al. Ten-dimensional anthropomorphic arm control in a human brainmachine interface: difficulties, solutions, and limitations. Journal of Neural Engineering 12, 016011 (2014).

7. Aflalo, T. et al. Decoding motor imagery from the posterior parietal cortex of a tetraplegic human. Science 348, 906–910 (2015).

8. Shannon, C. E. Prediction and Entropy of Printed English. Bell system technical journal 30, 50–64 (1951).

9. Karpathy, A., Johnson, J. & Fei-Fei, L. Visualizing and Understanding Recurrent Networks. arXiv preprint 1506.02078 (2015).

10. Radford, A. et al. Language Models are Unsupervised Multitask Learners. OpenAI blog 1, 9 (2019).

11. Gilja, V. et al. A high-performance neural prosthesis enabled by control algorithm design. Nature Neuroscience 15, 1752–1757 (2012).

12. Dangi, S., Orsborn, A. L., Moorman, H. G. & Carmena, J. M. Design and Analysis of Closed-Loop Decoder Adaptation Algorithms for Brain-Machine Interfaces. Neural Computation 25, 1693–1731 (2013).

13. Orsborn, A. L. et al. Closed-Loop Decoder Adaptation Shapes Neural Plasticity for Skillful Neuroprosthetic Control. Neuron 82, 1380–1393 (2014).

14. Silversmith, D. B. et al. Plug-and-play control of a brain–computer interface through neural map stabilization. Nature Biotechnology 39, 326–335 (2021).

15. Kim, S.-P., Simeral, J. D., Hochberg, L. R., Donoghue, J. P. & Black, M. J. Neural control of computer cursor velocity by decoding motor cortical spiking activity in humans with tetraplegia. Journal of Neural Engineering 5, 455 (2008).

16. Sussillo, D. et al. A recurrent neural network for closed-loop intracortical brain–machine interface decoders. Journal of Neural Engineering 9, 026027 (2012).

17. Sussillo, D., Stavisky, S. D., Kao, J. C., Ryu, S. I. & Shenoy, K. V. Making brain–machine interfaces robust to future neural variability. Nature Communications 7, 13749 (2016).

18. Kao, J. C. et al. Single-trial dynamics of motor cortex and their applications to brain-machine interfaces. Nature Communications 6, 7759 (2015).

19. Kao, J. C., Nuyujukian, P., Ryu, S. I. & Shenoy, K. V. A High-Performance Neural Prosthesis Incorporating Discrete State Selection With Hidden Markov Models. IEEE Transactions on Biomedical Engineering 64, 935–945 (2016).

20. Shenoy, K. V. & Carmena, J. M. Combining Decoder Design and Neural Adaptation in Brain-Machine Interfaces. Neuron 84, 665–680 (2014).

21. Lawhern, V. J. et al. EEGNet: a compact convolutional neural network for EEG-based brain–computer interfaces. Journal of Neural Engineering 15, 056013 (2018).

22. Forenzo, D., Zhu, H., Shanahan, J., Lim, J. & He, B. Continuous tracking using deep learning-based decoding for noninvasive brain–computer interface. PNAS Nexus 3, pgae145 (2024).

23. Shenoy, K. V., Sahani, M. & Churchland, M. M. Cortical Control of Arm Movements: A Dynamical Systems Perspective. Annual Review of Neuroscience 36, 337–359 (2013).

24. Fan, J. M. et al. Intention estimation in brain–machine interfaces. Journal of Neural Engineering 11, 016004 (2014).

25. Ganguly, K. & Carmena, J. M. Emergence of a Stable Cortical Map for Neuroprosthetic Control. PLoS Biology 7, e1000153 (2009).

26. Ganguly, K., Dimitrov, D. F., Wallis, J. D. & Carmena, J. M. Reversible large-scale modification of cortical networks during neuroprosthetic control. Nature Neuroscience 14, 662–667 (2011).

27. Pfurtscheller, G. & Da Silva, F. L. Event-related EEG/MEG synchronization and desynchronization: basic principles. Clinical Neurophysiology 110, 1842–1857 (1999).

28. Olsen, S. et al. An artificial intelligence that increases simulated brain–computer interface performance. Journal of Neural Engineering 18, 046053 (2021).

29. Schulman, J., Wolski, F., Dhariwal, P., Radford, A. & Klimov, O. Proximal Policy Optimization Algorithms. arXiv preprint 1707.06347 (2017).

30. Liu, S. et al. Grounding DINO: Marrying DINO with Grounded Pre-Training for Open-Set Object Detection. arXiv preprint 2303.05499 (2023).

31. Stieger, J. R. et al. Mindfulness Improves Brain–Computer Interface Performance by Increasing Control Over Neural Activity in the Alpha Band. Cerebral Cortex 31, 426–438 (2021).

32. Stieger, J. R., Engel, S. A. & He, B. Continuous sensorimotor rhythm based brain computer interface learning in a large population. Scientific Data 8, 98 (2021).

33. Edelman, B. J., Baxter, B. & He, B. EEG Source Imaging Enhances the Decoding of Complex Right-Hand Motor Imagery Tasks. IEEE Transactions on Biomedical Engineering 63, 4–14 (2016).

34. Scherer, R. et al. Individually adapted imagery improves brain-computer interface performance in end-users with disability. PloS one 10, e0123727 (2015).

35. Millan, J. d. R. et al. A local neural classifier for the recognition of EEG patterns associated to mental tasks. IEEE transactions on neural networks 13, 678–686 (2002).

36. Huang, D. et al. Decoding Subject-Driven Cognitive States from EEG Signals for Cognitive Brain– Computer Interface. Brain Sciences 14, 498 (2024).

37. Meng, J. et al. Noninvasive Electroencephalogram Based Control of a Robotic Arm for Reach and Grasp Tasks. Scientific Reports 6, 38565 (2016).

38. Reddy, S., Dragan, A. D. & Levine, S. Shared Autonomy via Deep Reinforcement Learning. arXiv preprint 1802.01744 (2018).

39. Laghi, M. et al. Shared-Autonomy Control for Intuitive Bimanual Tele-Manipulation. In 2018 IEEE-RAS 18th International Conference on Humanoid Robots (Humanoids), 1–9 (2018).

40. Tan, W. et al. On Optimizing Interventions in Shared Autonomy. In Proceedings of the AAAI Conference on Artificial Intelligence, vol. 36, 5341–5349 (2022).

41. Peng, Z., Mo, W., Duan, C., Li, Q. & Zhou, B. Learning from Active Human Involvement through Proxy Value Propagation. Advances in Neural Information Processing Systems (2023).

42. Yoneda, T., Sun, L., Stadie, B., Walter, M. et al. To the Noise and Back: Diffusion for Shared Autonomy. arXiv preprint 2302.12244 (2023).

43. McMahan, B. J., Peng, Z., Zhou, B. & Kao, J. C. Shared Autonomy with IDA: Interventional Diffusion Assistance. Advances in Neural Information Processing Systems (2024).

44. Brohan, A. et al. RT-1: Robotics Transformer for Real-World Control at Scale. arXiv preprint 2212.06817 (2022).

45. Brohan, A. et al. RT-2: Vision-language-action models transfer web knowledge to robotic control. arXiv preprint 2307.15818 (2023).

46. Nair, S., Rajeswaran, A., Kumar, V., Finn, C. & Gupta, A. R3M: A Universal Visual Representation for Robot Manipulation. arXiv preprint 2203.12601 (2022).

47. Ma, Y. J. et al. VIP: Towards Universal Visual Reward and Representation via Value-Implicit Pre-Training. arXiv preprint 2210.00030 (2022).

48. Willett, F. R. et al. A high-performance speech neuroprosthesis. Nature 620, 1031–1036 (2023).

49. Leonard, M. K. et al. Large-scale single-neuron speech sound encoding across the depth of human cortex. Nature 626, 593–602 (2024).

50. Card, N. S. et al. An accurate and rapidly calibrating speech neuroprosthesis. N. Engl. J. Med. 391, 609–618 (2024).

51. Sato, M. et al. Scaling law in neural data: Non-invasive speech decoding with 175 hours of eeg data. arXiv preprint 2407.07595 (2024).

52. CTRL-labs at Reality Labs, Sussillo, D., Kaifosh, P. & Reardon, T. A generic noninvasive neuromotor interface for human-computer interaction. bioRxiv 2024.02.23.581779 (2024).

53. Kao, J. C., Nuyujukian, P., Ryu, S. I. & Shenoy, K. V. A High-Performance Neural Prosthesis Incor-porating Discrete State Selection With Hidden Markov Models. IEEE Transactions on Biomedical Engineering 64, 935–945 (2017).

54. Nuyujukian, P. et al. Monkey models for brain-machine interfaces: The need for maintaining diversity. In Proceedings of the 33rd Annual Conference of the IEEE EMBS, vol. 2011, 1301–1305 (IEEE, Boston, Massachusetts, 2011).

55. Suminski, A. J., Tkach, D. C., Fagg, A. H. & Hatsopoulos, N. G. Incorporating Feedback from Multiple Sensory Modalities Enhances Brain-Machine Interface Control. Journal of Neuroscience 30, 16777–16787 (2010).

56. Kaufman, M. T., Churchland, M. M., Ryu, S. I. & Shenoy, K. V. Cortical activity in the null space: permitting preparation without movement. Nature Neuroscience 17, 440–448 (2014).

57. Kaufman, M. T. et al. The Largest Response Component in Motor Cortex Reflects Movement Timing but Not Movement Type. eNeuro 3, ENEURO.0085–16.2016 (2016).

58. Dangi, S. et al. Continuous closed-loop decoder adaptation with a recursive maximum likelihood algorithm allows for rapid performance acquisition in brain-machine interfaces. Neural Computation 26, 1811–1839 (2014).

59. Fitts, P. M. The information capacity of the human motor system in controlling the amplitude of movement. Journal of experimental psychology 47, 381 (1954).

60. Gramfort, A. et al. MNE software for processing MEG and EEG data. NeuroImage 86, 446–460 (2014).

