## Extended Data for "Non-invasive brain-machine interface control with artificial intelligence copilots"

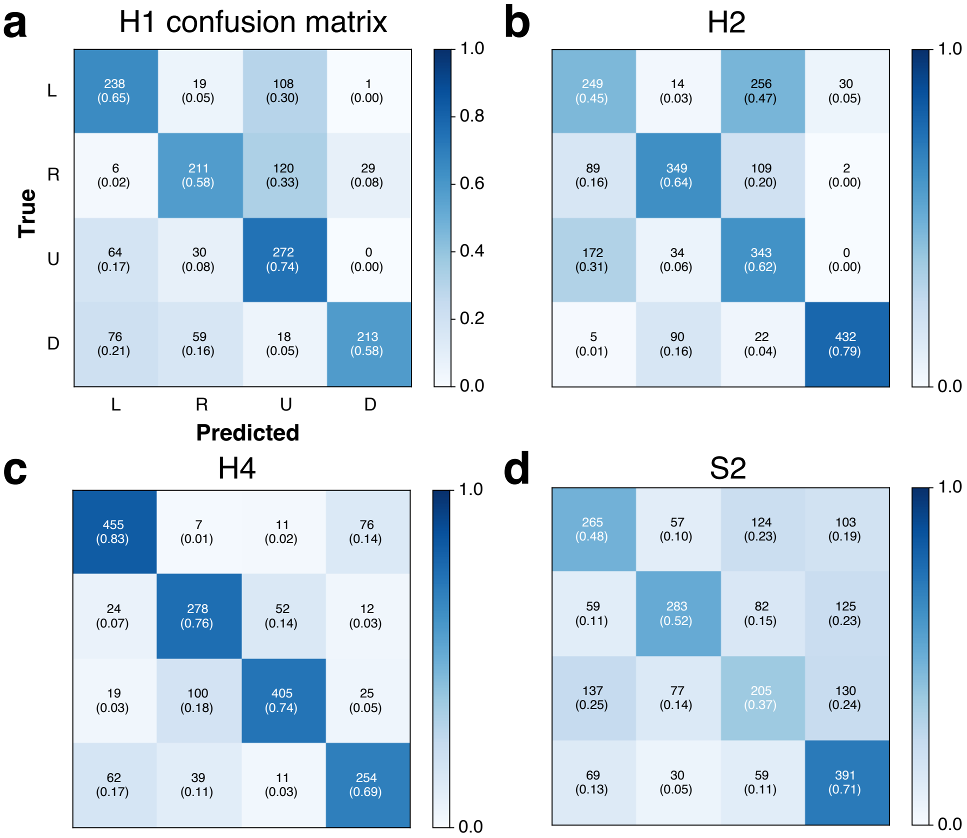


**Extended Data Figure 1.** Confusion matrices for the seed decoder from the open loop session that was used in the decorrelated session for **a-c,** healthy (H1,H2,H4) and **d,** SCI (S2) participants. All results are computed on test set data.

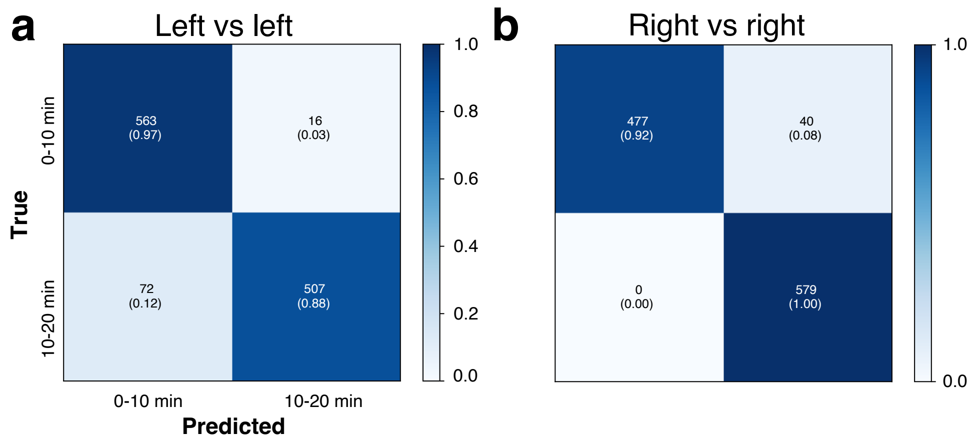


**Extended Data Figure 2.** **a,** Confusion matrix for decoding the left class in the first half (0-10 min) or second half (10-20 min) of the open loop session in test data. Although the motor intent is the same, EEG activity can be easily decoded from the first vs second half of the session, indicating decodable shifts in EEG activity.

12

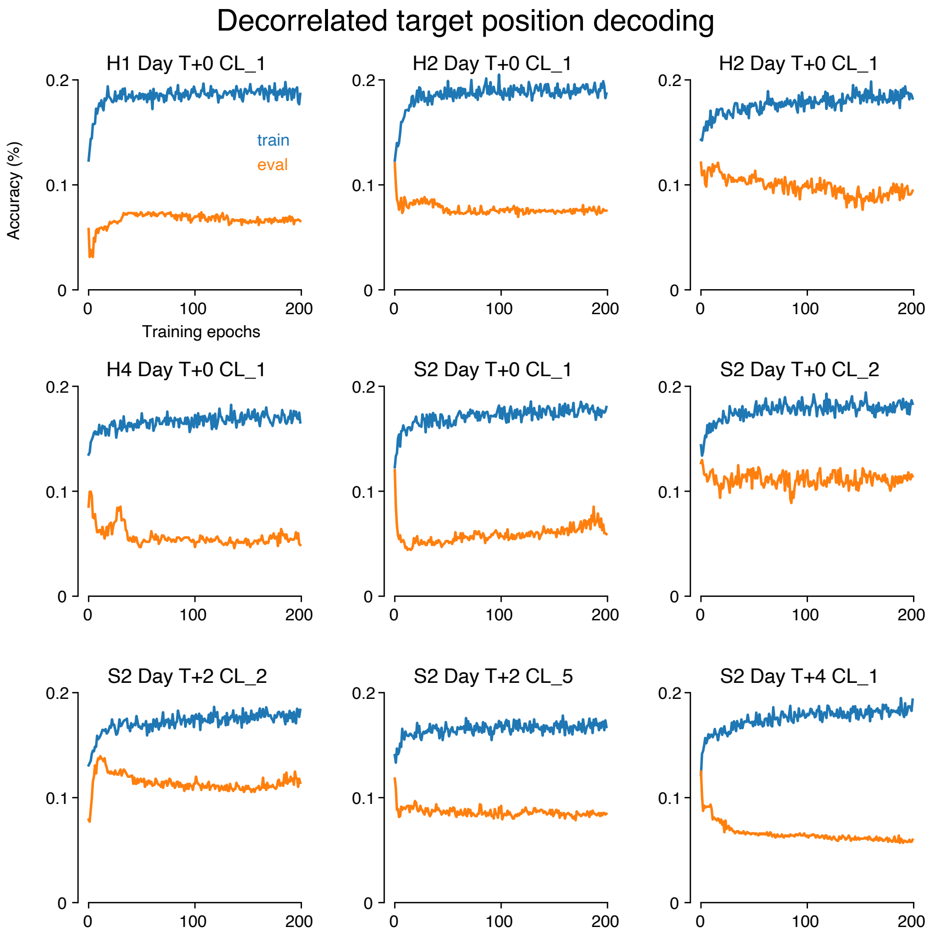


**Extended Data Figure 3.** Training accuracy (blue) and validation accuracy (orange) for decoding *target position* from the CNN output hidden state in all decorrelated training sets. Note that we recalibrated S2's CNN on two separate days (labeled T+2, T+4), which is why S2 has multiple decorrelated sessions. Eye movements and motor actions are decoupled in the decorrelated task, meaning that CNN features that decode motor intent should not to decode target position (e.g., what target the user is either looking at, or following the cursor to) above chance. This reflects that eye movement artifacts are not represented in the CNN hidden state. Consistent with this, we did not observe consistent above chance (12.5%) decoding target location from the CNN hidden state. After training, we often achieved worse than chance, indicating overfitting. We therefore show the validation accuracies across all epochs of training to demonstrate that validation accuracy does not achieve above chance accuracies throughout training.

13

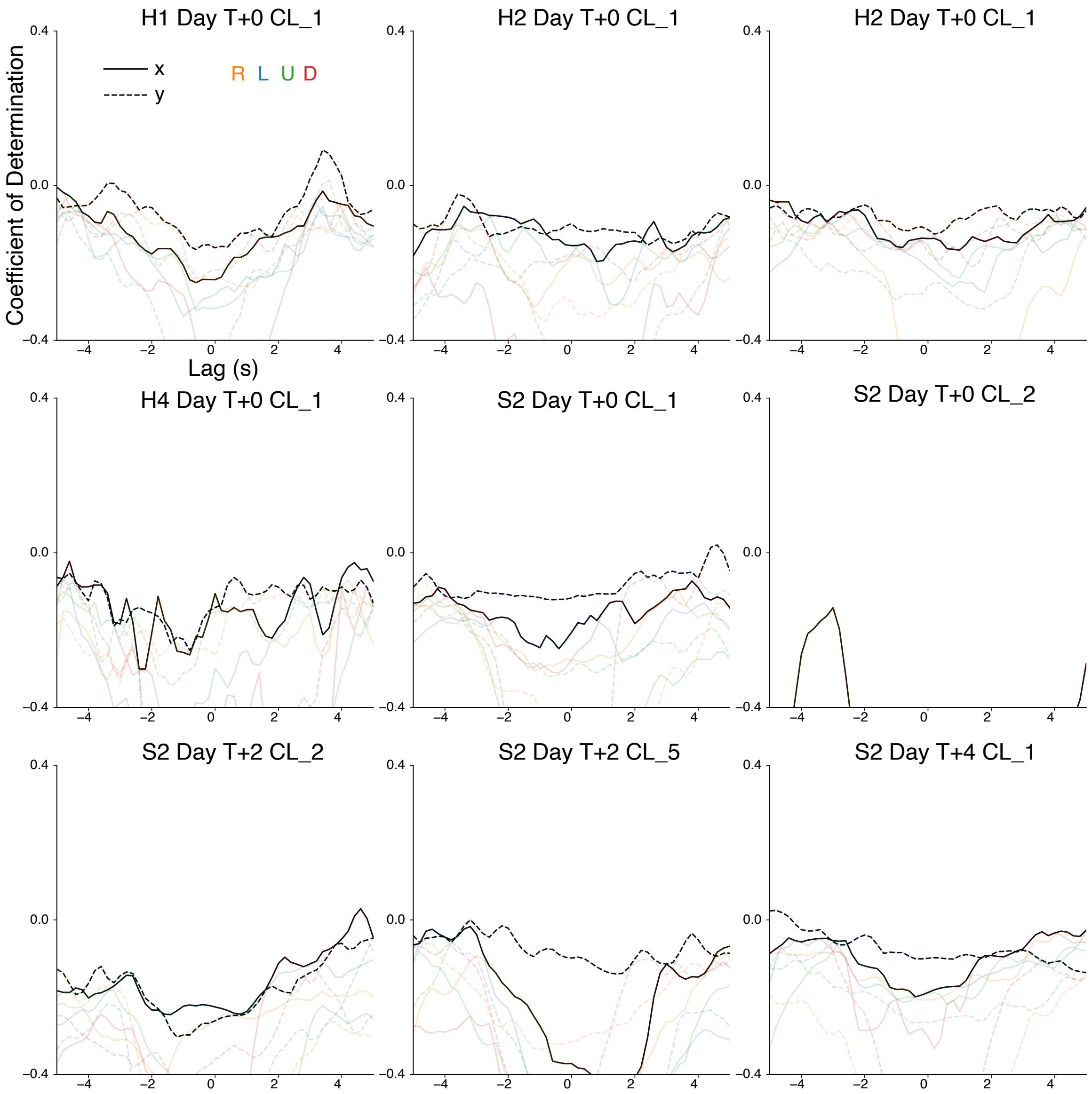


**Extended Data Figure 4.** Prediction of eye tracker gaze location from decoder hidden state during the decorrelated closed-loop task for a range of temporal offsets. Data for each session were split into 5 folds without shuffling. We trained linear regression models to predict gaze location from decoder hidden state, with one model trained from data for each motor prompt (right, left, up, down) for each offset for each data split. Lines show the maximum average coefficient of determination, by averaging each value across all 5 folds, then taking the maximum over all prompts. A delay of zero means that gaze data and hidden state are aligned as recorded, and a positive delay means that the gaze data at a certain timestamp is aligned with the hidden state recorded at a future timestamp. Black lines show the maximum average values, with individual averages shown as lighter colored lines. In general, coefficient of determinations were negative, meaning that these regression models could not predict the eye gaze data better than the mean on validation data. This provides strong evidence that the hidden state did not contain decodable eye movement signals.

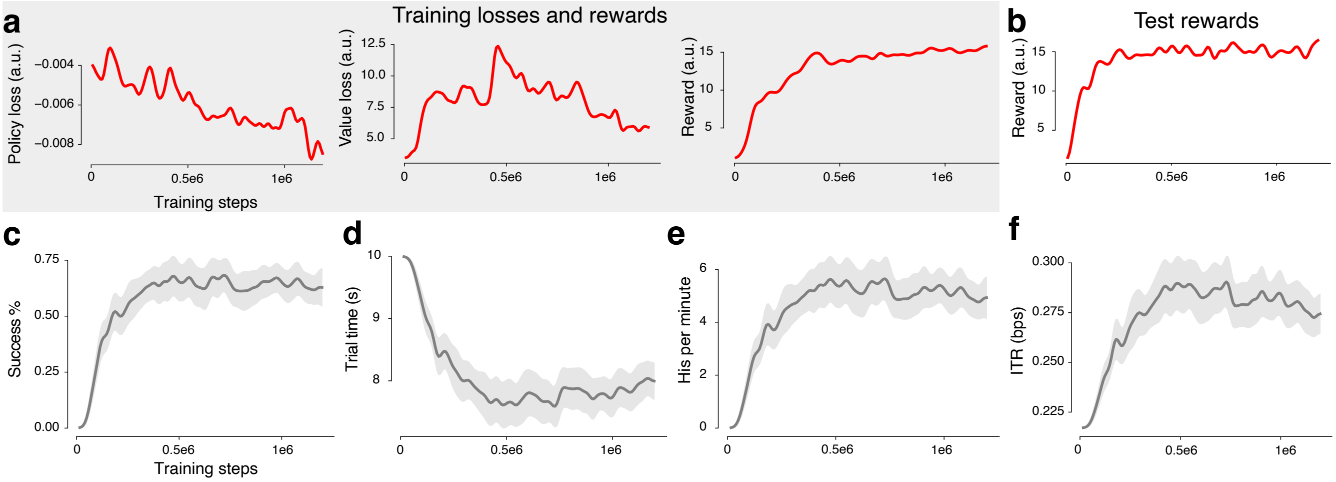


**Extended Data Figure 5.** **a,** PPO policy loss, value function loss, and rewards over training for the cursor copilot. The copilot increases rewards through training, as well as **b,** in an evaluation test environment, where the copilot was frozen every 8, 192 training steps and evaluated on the center-out and back task. **c,** Success percentage, **d,** trial time, **e,** target hit rate, and **f,** Fitts ITR on the center-out 8 task over the course of training. These demonstrate that the copilot learns to use the surrogate KF signals to perform the center-out 8 task. Please note that these numbers are in general lower (e.g., success percentage does not reach 100%) because the copilot task was more challenging, having a 2 second target hold time to encourage goal acquisition behavior (see Methods).

15

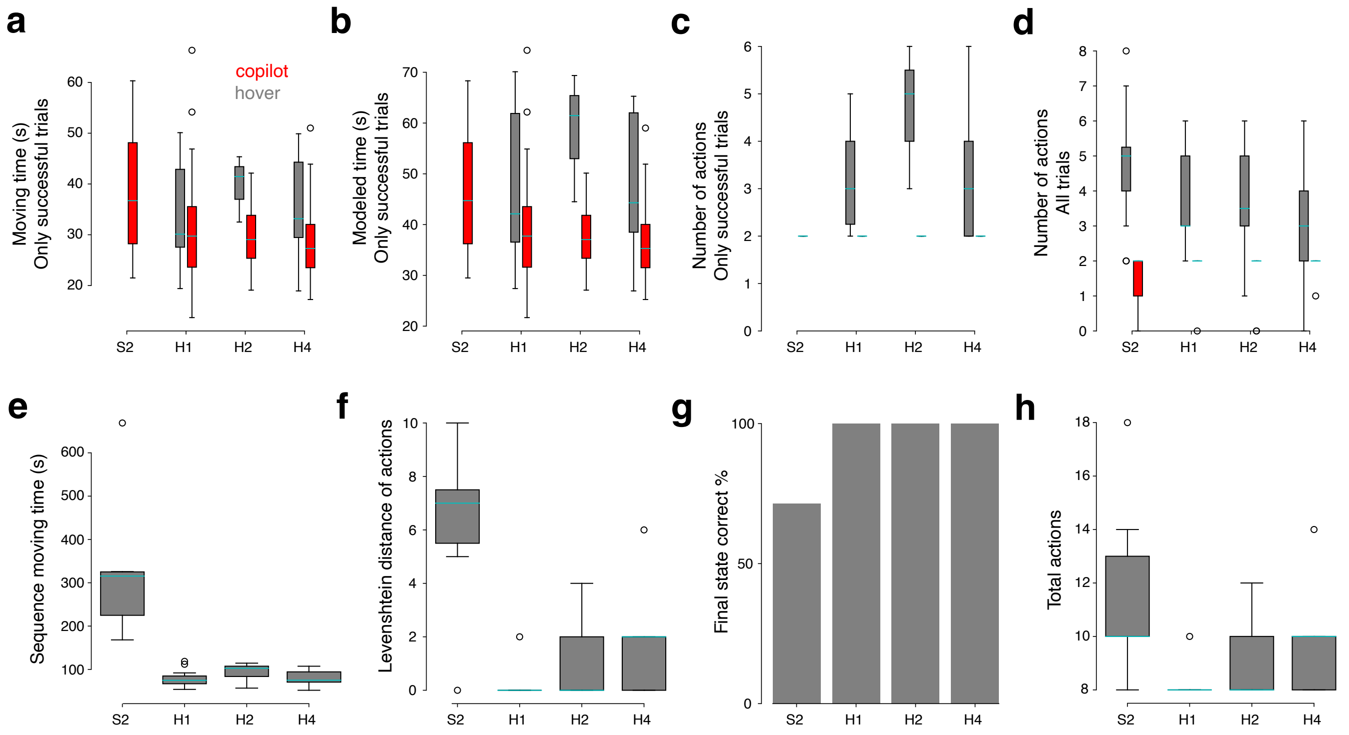


**Extended Data Figure 6.** **a,** Moving time (excluding time required for raising/lowering arm and closing/opening the gripper) for the Paired Pick-And-Place task for only successful trials. This moving time is a conservative estimate of the copilot improvement, since the copilot helps to successfully grasp and place blocks. In this example, any time spent trying to pick and place blocks without the copilot is excluded, since this only includes time spent moving towards pick and place locations. Note, S2 was unable to perform any trials successfully using only hover, which is why there is no data for the hover condition for S2. **b,** This estimates the total trial time in the Paired Pick-And-Place task if every action took 4 seconds to attempt or complete. This estimate, in general, underestimates the actual trial time for hover trials. Nevertheless, the copilot still improves overall time spent to perform the task. **c,** Actions are attempted pick and place attempts on eventually successful trials. For the copilot, all successful trials had one successful pick and one successful place. For hover trials, the median number of actions in successful trials was generally higher due to failed pick and place attempts. **d,** The same as **c**, but including failed trials. Note that the number of actions can be less than 2 if zero or one attempted action was made throughout the trial. **e,** Distribution of moving time for the sequential pick-and-place task. **f,** Levenshtein edit distance between prompted action sequences and actual action sequences. The Levenshtein edit distance measures the total number of insertions, deletions, and substitutions required to change one sequence to another. In this instance, it is the minimum number of actions in the user's performed action sequence that must be inserted, deleted, or substituted to produce the prompted action sequence. **g,** Percentage of sequences in which users placed all blocks on the correct target at the end of the trial. Even though there were errors throughout each trial (e.g., placing a block on the wrong cross), these errors could be corrected (e.g., picking up the block from the wrong cross and placing it on the correct cross). **h,** Total number of actions (adjusted to include missing actions for incomplete trials) for all sequence task trials. The minimum number of actions to successfully pick and place 4 blocks at the correct location is 8.

| | 1 | 2 | 3 | 4 | 5 |
|---|---|---|---|---|---|
| S2 | 2024-02-12 | 2024-02-13 | 2024-02-21 | 2024-02-27 | 2024-03-18 |
| H1 | 2024-02-13 | 2024-02-14 | 2024-02-15 | 2024-02-20 | 2024-02-21 |
| H2 | 2024-02-02 | 2024-02-05 | 2024-02-06 | 2024-02-07 | 2024-02-09 |
| H4 | 2024-03-01 | 2024-03-04 | 2024-03-06 | 2024-03-11 | 2023-03-13 |

**Extended Table 1.** Dates on which the experiments were performed to collect data from the participants.

| Parameter | Description | Value |
|---|---|---|
| # Temporal Filters | Number of temporal filters in the first convolutional layer | 8 |
| Temporal Filter Size | Size of each temporal filter (time × frequency) | (1, 51) |
| # Spatial Filters | Number of spatial filters in the spatial convolutional layer | 2 |
| Spatial Filter Size | Size of each spatial filter (channels × 1) | (58, 1) |
| Average Pool Size (1st) | Size of the average pool filter in the first block | (1, 3) |
| Separable Conv Size | Size of the separable convolution in the second block | (1, 16) |
| Average Pool Size (2nd) | Size of the average pool filter in the second block | (1, 16) |
| Dropout | Dropout rate applied for both layers | 0.5 |
| Learning Rate | Initial learning rate | 0.001 |
| Learning Rate Schedule | Reduction factor when validation accuracy does not improve | 0.1 |
| Weight Decay | Weight decay applied during optimization | 0.0001 |
| Batch Size | Number of samples in each batch | 32 |

**Extended Table 2.** Detailed parameters of EEGNet.

Reward and Action Hyperparameter

| Parameter | Description | Value |
|---|---|---|
| $\alpha_{\text{KF}}$ | Contribution of KF-CNN | 0.6 |
| $\alpha_{\text{CP}}$ | Contribution of RL copilot | 0.3 |
| $K$ | Electrostatic constant | 1 |
| $q$ | Fixed electric charge on cursor | 1 |
| $\varepsilon$ | Epsilon stabilization in the electrostatic equation for avoiding division by zero | 0.01 |
| $k$ | Linear reward scale | 66.67 |
| $\gamma$ | Reward discount factor | 0.98901 |

Copilot Training Hyperparameter

| Parameter | Description | Value |
|---|---|---|
| learning rate | Copilot learning rate | 0.0003 |
| rollout buffer | Number of steps stored before gradient update | 2048 |
| batch size | Mini batch size | 512 |
| total learning steps | Total number of steps taken for training | 1,200,000 |
| delay | Delay used in $p_{\text{surrogate}}(\mathbf{v}_t^{\text{KF}})$ | $U[300ms, 600ms]$ |
| $\sigma$ | Variance of angular noise in $p_{\text{surrogate}}(\mathbf{v}_t^{\text{KF}})$ | $\pi/3$ |
| $\Sigma$ | Variance of additive Gaussian noise in $p_{\text{surrogate}}(\mathbf{v}_t^{\text{KF}})$ | $0.03\mathbf{I}$ |

**Extended Table 3.** Detailed parameters of RL cursor copilot.
